# A prospective multi-cohort study identifies reproducible molecular responses to musical stimulation in the human tear proteome

**DOI:** 10.64898/2026.07.31.737062

**Authors:** Alba Camino-Mera, María José Currás-Tuala, Nour El Zahraa Mallah, Jacobo Pardo-Seco, Conrado Martínez-Cadenas, Laura Navarro, Carmen Pena, Susana B. Bravo, Federico Martinón-Torres, Alberto Gómez-Carballa, Antonio Salas

## Abstract

Music elicits complex emotional, cognitive, and physiological responses, yet the molecular mechanisms underlying these effects remain poorly understood. Tear fluid is an accessible biofluid that reflects both systemic and local physiology, providing a unique opportunity to investigate the biological effects of musical stimulation. Quantitative mass spectrometry-based proteomics was performed on paired tear samples (n = 336) collected before and after exposure to a standardized musical stimulus in two independent cohorts of healthy participants, including both music-stimulated and music-unstimulated individuals. Integrative analyses, including differential expression, functional enrichment, co-expression network, correlation, and machine-learning approaches, identified reproducible molecular responses across cohorts. Fourteen proteins, including PRTN3, LEG3, CATD, IGHG2, and ZA2G, remained significant after multiple-testing correction and showed concordant regulation in the stimulated cohorts but not in controls. Functional enrichment converged on innate immune, inflammatory, host-defense, and extracellular signaling pathways, and consensus network analysis identified reproducible immune-related modules. The 14-protein signature robustly discriminated pre- and post-stimulation samples, with no effect detected in the unstimulated control groups. These findings support tear proteomics as a promising non-invasive approach for investigating the molecular basis of auditory and emotional experiences.

## Introduction

Music is a universal form of human expression that transcends cultural and geographical boundaries. Beyond its artistic and social significance, it influences emotional, cognitive, and physiological processes. Increasing evidence shows that music modulates neural networks, promotes neuroplasticity, and affects behavioral and cognitive outcomes in both healthy individuals and patients with neurodegenerative diseases ^1–3^. In healthy individuals, musical engagement has been associated with improved cognitive performance and may help counteract aspects of cognitive aging ^4, 5^. Music-based interventions have also shown beneficial effects in neurodegenerative diseases, including Alzheimer’s disease, where they have been linked to improvements in memory, mood, autobiographical recall, and other cognitive and behavioral functions ^3, 6^.

Although the behavioral and clinical effects of music have been widely investigated, the underlying biological mechanisms remain poorly understood. Recent advances in omics technologies have revealed measurable molecular changes associated with music exposure in both healthy individuals and patients with neurodegenerative disorders ^7–13^. Genomics, metagenomics, and transcriptomics studies have identified alterations in gene expression, microbial communities, and biological pathways. Recent work further demonstrated music-associated molecular changes in blood ^7^ and saliva ^10^, supporting the hypothesis that auditory stimulation elicits systemic biological responses.

Despite these advances, whether music exposure induces measurable changes in the human proteome remains unknown. This represents an important knowledge gap because proteins are the primary functional mediators of cellular activity and provide a more direct representation of physiological processes than gene expression alone. Proteomics, the large-scale study of protein abundance, function, modifications, and interactions ^14^, has become a powerful approach for investigating physiological stress, disease progression, and therapeutic interventions ^15, 16^. However, proteomic studies in neuroscience often rely on cerebrospinal fluid or brain tissue, which are invasive and impractical for large-scale or longitudinal studies ^17, 18^.

Tear fluid is an attractive alternative because it can be collected rapidly, non-invasively, and cost-effectively. In addition to reflecting ocular conditions, tears contain a complex mixture of proteins and other biomolecules that can provide information about systemic physiological and pathological processes ^19–21^. As tear composition reflects both local and systemic biology, it offers a unique opportunity to investigate subtle molecular responses induced by external sensory stimuli. Nevertheless, no studies have examined whether musical stimulation influences the human tear proteome.

This study investigated the effects of music exposure on the tear proteome of healthy individuals using a quantitative proteomics approach to identify proteins and biological pathways modulated by auditory stimulation. It was hypothesized that musical stimulation would induce measurable alterations in tear protein composition, providing new insights into the molecular mechanisms underlying the biological effects of music and further supporting tear fluid as a promising non-invasive biofluid for biomarker discovery.

## Methods

### Experimental design and sampling

Within the Sensogenomics project (http://sensogenomics.com) ^22^, we conducted a prospective controlled repeated-measures study comprising two independent music-stimulated cohorts and two concurrent music-unexposed control cohorts (**Figure 1**). The music-stimulated cohorts were designed as an independent discovery cohort (2024; *n*=88) and an independent replication cohort (2025; *n*=60) and were based on two independent 50-minute live classical music concerts held under comparable conditions at the Auditorio de Galicia (Santiago de Compostela, Spain). Participants were unaware of the musical repertoire, remained seated throughout the performances, and engaged exclusively in passive listening. Both concerts followed the same emotional framework, with a sadness-evoking first half and a joy-evoking second half. Two contemporaneous control cohorts (*n*=11 and *n*=9) remained in a separate sound-isolated room under the same temporal schedule and environmental conditions, without exposure to music or structured sensory stimulation. Tear samples were collected from all participants using Schirmer strips without anesthesia immediately before (TP1) and after (TP2) the concert or control session. Participants with dry eye disease or other ocular pathologies were excluded. Pooled samples were used to construct an in-house spectral library, and quantitative proteomic analyses were subsequently performed on individual paired samples to compare protein abundance between TP1 and TP2. The cohorts were broadly comparable in their demographic characteristics, with no significant differences in sex distribution across groups and only a single significant difference in age between the 2025 cohort and the 2024 control group (**Table S1**). A more detailed description is provided in **Supplementary Text 1**; see also **Figure 1**.

**Figure 1.**
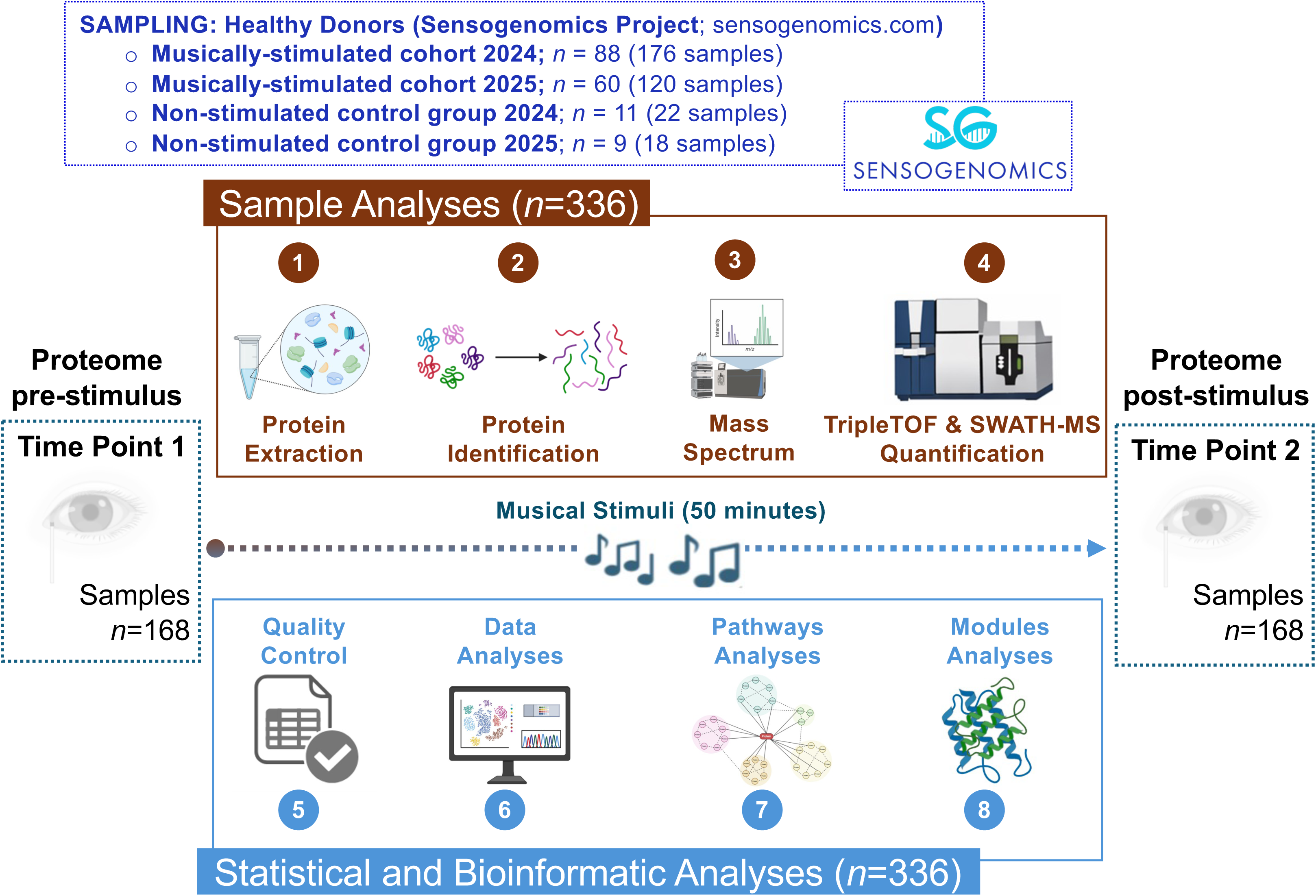
Schematic overview of the experimental and analytical workflow for the four sample groups included in the present study: the two music- unstimulated control groups (2024 and 2025) and the two independent music-stimulated cohorts (2024 and 2025).

### Tear preparation for protein analysis

Tear proteins were extracted by incubating the Schirmer strips in 100 μL of 100 mM ammonium bicarbonate at room temperature for 1 h, followed by centrifugation (20 min at 13,000×g) ^23, 24^. Then, the supernatant and precipitated proteins were collected using the MeOH/CHCl_3_ method. The protein concentration was determined using the RC-DC kit (Bio-Rad) ^23–25^.

For global protein identification, 100 μg of total protein was concentrated on a 10% SDS-PAGE gel, and electrophoresis was stopped once the dye front had migrated approximately 3 mm into the resolving gel ^26^. We visualized the proteins with Sypro Ruby staining (Lonza), excised them, and subjected them to in-gel tryptic digestion ^27^. We extracted the peptides by sequential incubations in 60% acetonitrile with 0.5% formic acid, pooled them, concentrated them using a SpeedVac, and stored them at −20 °C. To minimize batch effects during sample processing (tear preparation and subsequent technical analyses; see below), all TP1 and TP2 samples from the 2024 cohort, together with the control group samples, were processed in the same experimental batch, whereas all samples from the 2025 cohort were processed together in a separate batch.

### SWATH-MS protein quantification

A detailed description of the SWATH-MS workflow is provided in **Supplementary Text 1**. Briefly, pooled tear samples from each experimental group were analyzed by shotgun data-dependent acquisition (DDA) microLC-MS/MS using a TripleTOF 6600 mass spectrometer (SCIEX) to generate an in-house spectral library. Peptides and proteins were identified against the UniProt human database using ProteinPilot software, applying a false discovery rate (FDR) of <1% at both peptide and protein levels. The resulting high-confidence peptide spectra were used to construct the spectral library for subsequent SWATH data-independent acquisition (DIA) analysis.

Individual tear samples were then analyzed by SWATH-MS using the same chromatographic conditions. Variable precursor isolation windows were optimized according to precursor ion density, enabling comprehensive and reproducible protein quantification across all samples. Fragment-ion chromatograms were extracted and quantified in *PeakView* using the DDA-derived spectral library. Up to ten peptides per protein and seven fragment ions per peptide were used for quantification, with only peptides meeting an FDR of <1% retained. Protein abundances were calculated by summing peptide peak areas and globally normalized using the total signal intensity of each sample to ensure reliable comparison across experimental groups.

### Data processing and statistical analysis

To conduct statistical analyses and all data processing, an in-house pipeline was developed in Python (v3.9.6) and R (v4.5.2). Proteins with >30% missing values across samples were removed. Proteins exhibiting zero variance across individuals were also excluded from further analysis. Subsequently, data normalization was performed using Variance Stabilizing Normalization (VSN) implemented in the R package *vsn* ^28^.

For visualization analyses, including Principal Component Analysis (PCA) and heatmap generation, missing values were imputed using the median value of each protein. Subject-specific effects were further adjusted using the *removeBatchEffect* function from the R package *limma* ^29^, and PCA was performed on the adjusted data.

Differential protein expression analyses were conducted using *limma*. This framework implements linear models for microarray and high-throughput data analysis, applying moderated *t*-statistics based on an empirical Bayes approach ^30^. To account for repeated measurements within individuals, intra-subject correlation was estimated using the *duplicateCorrelation* function. Log_2_ fold changes (log_2_FC) and *P*-values were computed for each protein. Multiple testing correction was performed using the FDR method to control for false-positive findings.

Statistical analyses were performed independently for the 2024 and 2025 cohorts, including both stimulated and unstimulated donors. In addition, a combined analysis including samples from both cohorts was conducted to assess the reproducibility of the findings after increasing the sample size. For the combined dataset, batch effects associated with the year of sample collection were corrected using the ComBat function from the R package *sva* ^31^. The batch-corrected data were subsequently used for differential expression analysis.

Differentially expressed proteins (DEPs) were defined as those with a *P*_adj_<0.05. Protein–protein interaction (PPI) networks were generated in STRING (https://string-db.org) ^32^ using the DEPs identified in each cohort. Only interactions with a confidence score >0.4 were retained. Clustering was performed using the Markov Cluster Algorithm (MCL; inflation=3) to identify functionally related groups of interacting proteins and their associated biological processes.

Functional enrichment analyses were performed using *clusterProfiler* package ^33^, employing *org.Hs.eg.db* annotation database ^34^ for Gene Ontology (GO) Biological Process terms ^35^ and the Reactome PA package for Reactome pathway enrichment ^36^. GO Biological Process terms and Reactome pathways were performed separately for the 2024 and 2025 cohorts using the most significant DEPs, applying the criteria *P*_adj_<0.05 and |log_2_FC|≥0.5. We first investigated the top 20 most significantly enriched pathways in each cohort separately. Functional enrichment analyses were then repeated separately for the up- and down-regulated DEPs to determine the overall direction of pathway regulation. In a separate analysis, all significantly enriched pathways identified in the 2024 and 2025 cohorts were aggregated into broader functional categories based on shared lexical terms.

Volcano plots, heatmaps, PCA plots, and boxplots were generated using visualization packages in Python and R, including *matplotlib* ^37^, *seaborn* ^38^, *ggplot2* ^39^, and *ComplexHeatmap* ^40^.

### Weighted protein co-expression network analysis

Weighted protein co-expression network analysis was performed using the *WGCNA* R package ^41^. A consensus network was constructed from the 2024 and 2025 cohorts using the 203 proteins shared between both datasets to identify reproducible protein modules associated with musical stimulation. Normalized protein abundance data were corrected for repeated measures using the *removeBatchEffect* function from the *limma* package ^29^, and a signed weighted network was generated using a soft-thresholding power of β=5. Consensus modules were identified through hierarchical clustering and summarized using module eigengenes, which were tested independently in each cohort for association with stimulation status (TP1 *vs*. TP2). Module preservation, hub protein identification, and functional enrichment analyses were subsequently performed using *WGCNA* and *clusterProfiler* ^33^ with FDR correction. A detailed description of the network construction, preservation analysis, hub ranking, and enrichment procedures is provided in **Supplementary Text 1**.

### Classification performance of replicated proteins

To assess whether the replicated proteins captured musical stimulation-associated molecular differences, their ability to discriminate between TP1 and TP2 samples was assessed through the application of four different supervised machine-learning models implemented with the *scikit-learn* library ^42^: logistic regression (LR), random forest (RF), support vector machine (SVM), and linear discriminant analysis (LDA). VSN-normalized protein abundance values, normalized across samples rather than within individual patients, were used as predictors. Models were trained using the 2024 cohort and subsequently evaluated on the independent 2025 cohort. Default *scikit-learn* hyperparameters were used unless otherwise specified. This analysis assessed the classification performance of the replicated proteins for discriminating between stimulation states and was not intended to develop or validate a clinically applicable biomarker signature.

## Results

### Differential tear protein expression following musical stimulation

A total of 212 proteins were initially quantified, representing approximately 80% of the spectral library. After quality-control filtering, proteins with >30% missing values or zero variance were excluded. In total, 203 proteins were consistently quantified across all cohorts and were therefore retained for cross-cohort comparative analyses.

Cohort 2024 included 88 individuals (176 paired samples collected before [TP1] and after [TP2] the musical intervention). PCA based on the 96 DEPs showed clear separation between TP1 and TP2 along PC1 (40.7% variance explained; **Figure 2A**). Differential expression analysis identified 96 DEPs (*P*_adj_<0.05), including 48 upregulated and 48 downregulated proteins (**Table S2**; **Figure 2B–D**). Thirty-two proteins (33.3%) showed |log2FC|≥0.5, with the strongest changes observed for A1AT, APOA1, APOA2, CERU, DEF3, DMBT1, HBA, HBD, IGHG2, KV401, LYST, PROL1, and PROL4. Hierarchical clustering based on the top 30 DEPs largely separated TP1 and TP2 samples (**Figure 2E**).

**Figure 2.**
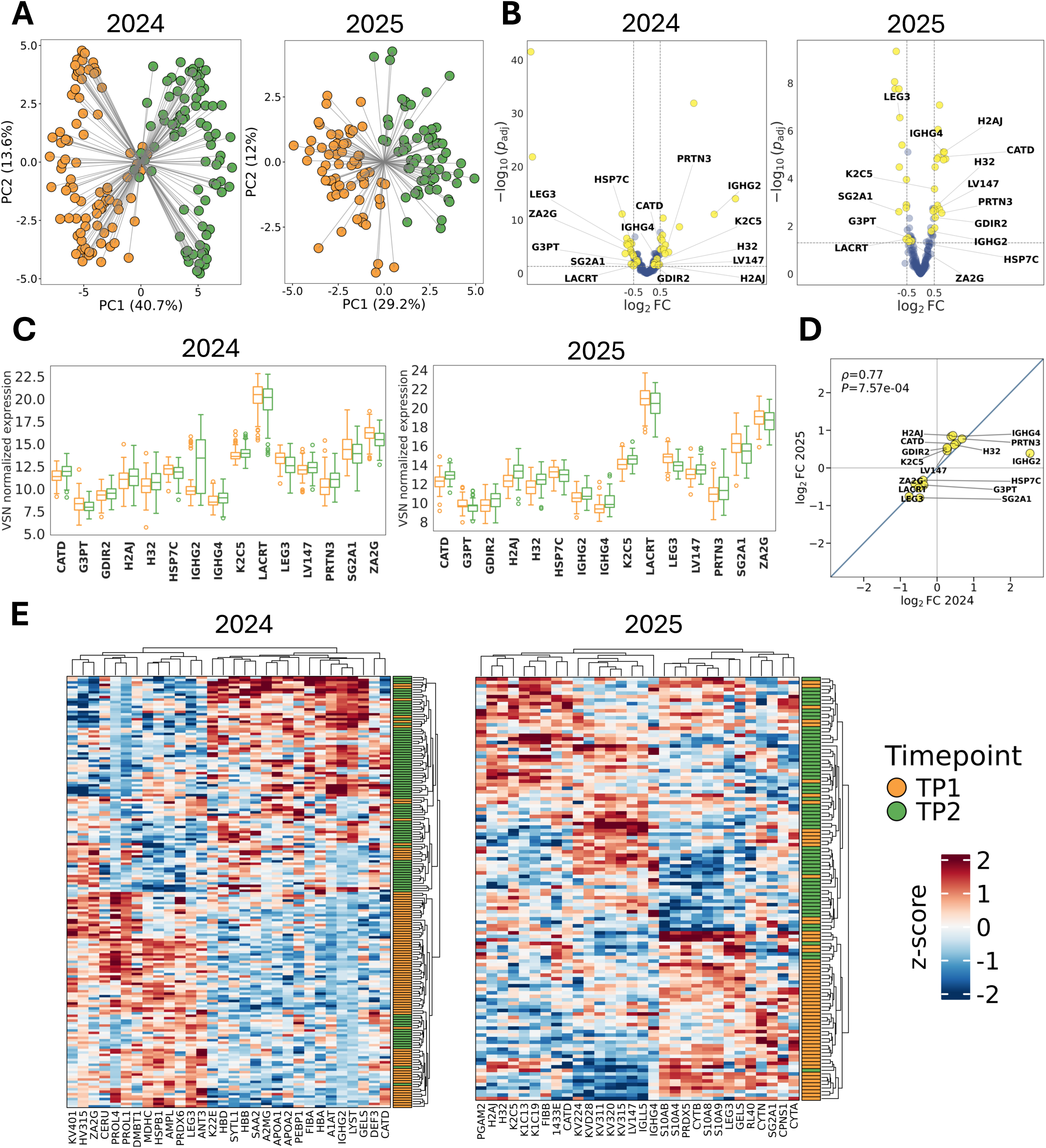
Differential protein expression analyses in the 2024 and 2025 music-stimulated cohorts following musical stimulation. (A) Principal component analyses (PCA) of samples collected before (TP1) and after (TP2) musical stimulation in the 2024 and 2025 cohorts, showing the overall separation between experimental conditions and based on the DEPs identified in each cohort. Arrows connect paired TP1 and TP2 samples from the same participant. (B) Volcano plots summarizing the differential expression analyses. The x-axis represents the log_2_ fold change (log_2_FC) between post- and pre-stimulation samples, whereas the y-axis represents the adjusted statistical significance (−log_10_ *P*_adj_) of the observed differences. Labeled proteins correspond to the 15 reproducible DEPs identified in both cohorts, which remained significant after multiple-testing correction and exhibited concordant directions of regulation. (C) Boxplots showing the distribution of normalized expression values for the 14 reproducible DEPs identified in both cohorts. (D) Scatter plots showing the correlation of log_2_FC values between the 2024 and 2025 cohorts for the 14 replicated proteins. (E) Heatmaps displaying the top 30 DEPs identified in the 2024 and 2025 cohorts, illustrating their expression patterns across samples and their capacity to discriminate between pre- and post-stimulation conditions.

Cohort 2025 included 60 participants (120 paired samples). PCA again showed clear TP1–TP2 separation along PC1 (29.2% variance explained; **Figure 2A**). Differential expression analysis identified 58 DEPs (*P*_adj_<0.05), including 31 upregulated and 27 downregulated proteins (**Table S3**; **Figure 2B–D**). Thirty proteins (51.7%) showed |log_2_FC|≥0.5, with the strongest changes observed for CATD, CYTB, GELS, H2AJ, IGHG4, K1C19, K2C5, KV311, KV315, KV320, LEG3, S10A4, S10A8, S10A9, S10AB, and FIBB. Clustering of the top 30 DEPs also separated TP1 and TP2 samples **(Figure 2E)**.

Both cohorts showed a similar balance of up- and down-regulated proteins. Although expression changes were larger in the 2024 cohort (log_2_FC: −4.37 to 3.33) than in the 2025 cohort (−0.95 to 0.92), the overall proteomic response was highly consistent (**Figure 2B–D**).

Cross-cohort comparison identified 26 proteins significant (*P*_adj_<0.05) in both cohorts, of which 15 showed concordant regulation and were considered replicated (**Table 1**; **Figure 2C**): LACRT, LEG3, IGHG4, H2AJ, CATD, K2C5, H32, LV147, SG2A1, PRTN3, GDIR2, IGHG2, G3PT, HSP7C, and ZA2G. These included nine upregulated and six downregulated proteins involved in innate immunity, immunoglobulin responses, epithelial homeostasis, lacrimal physiology, and inflammatory regulation. Log_2_FC values were strongly correlated between cohorts (Spearman’s ρ=0.77, *P*=7.57×10^−04^; **Figure 2D**), whereas neither music cohort correlated with the unstimulated control groups (**Figure S1**). Moreover, analysis of the combined 2024 and 2025 datasets further increased the statistical significance of the replicated proteins (**Table 1**).

**Table 1.** Fourteen-protein signature comprising differentially expressed proteins (*P*_adj_<0.05) reproducibly identified in both the 2024 and 2025 music-stimulated cohorts, with concordant directions of regulation.

| Protein | Gene | UniProt | 2024 cohort |  |  | 2025 cohort |  |  | 2024 + 2025 cohort |  |  |
| --- | --- | --- | --- | --- | --- | --- | --- | --- | --- | --- | --- |
|  |  |  | Log<sub>2</sub>FC | P-value | P<sub>adj</sub> | Log<sub>2</sub>FC | P-value | P<sub>adj</sub> | Log<sub>2</sub>FC | P-value | P<sub>adj</sub> |
| IGHG2 | *IGHG2* | P01859 | 2.53 | $2.5 \times 10^{-13}$ | $8.5 \times 10^{-12}$ | 0.39 | $3.20 \times 10^{-03}$ | $1.63 \times 10^{-02}$ | 1.3 | $3.2 \times 10^{-10}$ | $1.1 \times 10^{-08}$ |
| LEG3 | *LGALS3* | P17931 | -0.76 | $2.5 \times 10^{-07}$ | $3.1 \times 10^{-06}$ | -0.76 | $8.30 \times 10^{-09}$ | $2.81 \times 10^{-07}$ | -0.78 | $7.5 \times 10^{-14}$ | $3.9 \times 10^{-12}$ |
| CATD | *CTSD* | P07339 | 0.51 | $3.8 \times 10^{-06}$ | $3.6 \times 10^{-05}$ | 0.65 | $7.60 \times 10^{-07}$ | $1.29 \times 10^{-05}$ | 0.55 | $1.3 \times 10^{-10}$ | $5.3 \times 10^{-09}$ |
| ZA2G | *AZGP1* | P25311 | -0.61 | $1.5 \times 10^{-05}$ | $1.2 \times 10^{-04}$ | -0.43 | $1.21 \times 10^{-02}$ | $4.47 \times 10^{-02}$ | -0.56 | $8.2 \times 10^{-07}$ | $1.2 \times 10^{-05}$ |
| PRTN3 | *PRTN3* | P24158 | 0.68 | $3.1 \times 10^{-05}$ | $2.0 \times 10^{-04}$ | 0.77 | $4.31 \times 10^{-04}$ | $2.82 \times 10^{-03}$ | 0.69 | $2.7 \times 10^{-07}$ | $5.0 \times 10^{-06}$ |
| IGHG4 | *IGHG4* | P01861 | 0.38 | $4.4 \times 10^{-04}$ | $1.8 \times 10^{-03}$ | 0.83 | $3.96 \times 10^{-07}$ | $7.78 \times 10^{-06}$ | 0.56 | $6.1 \times 10^{-09}$ | $1.6 \times 10^{-07}$ |
| HSP7C | *HSPA8* | P11142 | -0.38 | $9.6 \times 10^{-04}$ | $3.5 \times 10^{-03}$ | -0.32 | $1.11 \times 10^{-02}$ | $4.17 \times 10^{-02}$ | -0.38 | $1.2 \times 10^{-05}$ | $1.2 \times 10^{-04}$ |
| G3PT | *GAPDHS* | O14556 | -0.36 | $2.4 \times 10^{-03}$ | $7.6 \times 10^{-03}$ | -0.45 | $6.96 \times 10^{-03}$ | $3.00 \times 10^{-02}$ | -0.39 | $2.3 \times 10^{-04}$ | $1.1 \times 10^{-03}$ |
| K2C5 | *KRT5* | P13647 | 0.28 | $4.0 \times 10^{-03}$ | $1.2 \times 10^{-02}$ | 0.51 | $2.59 \times 10^{-05}$ | $2.76 \times 10^{-04}$ | 0.34 | $6.5 \times 10^{-06}$ | $7.9 \times 10^{-05}$ |
| H32 | *HIST2H3A* | Q71DI3 | 0.47 | $7.9 \times 10^{-03}$ | $2.1 \times 10^{-02}$ | 0.62 | $1.38 \times 10^{-04}$ | $1.33 \times 10^{-03}$ | 0.51 | $4.8 \times 10^{-05}$ | $3.1 \times 10^{-04}$ |
| LV147 | *IGLV1-47* | P01700 | 0.27 | $9.5 \times 10^{-03}$ | $2.5 \times 10^{-02}$ | 0.46 | $3.32 \times 10^{-04}$ | $2.33 \times 10^{-03}$ | 0.32 | $1.1 \times 10^{-04}$ | $6.3 \times 10^{-04}$ |
| SG2A1 | *SCGB2A1* | O75556 | -0.47 | $1.0 \times 10^{-02}$ | $2.6 \times 10^{-02}$ | -0.79 | $3.66 \times 10^{-04}$ | $2.48 \times 10^{-03}$ | -0.63 | $1.7 \times 10^{-05}$ | $1.5 \times 10^{-04}$ |
| GDIR2 | *ARHGDIB* | P52566 | 0.27 | $1.1 \times 10^{-02}$ | $2.7 \times 10^{-02}$ | 0.54 | $6.77 \times 10^{-04}$ | $4.29 \times 10^{-03}$ | 0.35 | $1.1 \times 10^{-04}$ | $6.3 \times 10^{-04}$ |
| H2AJ | *H2AFJ* | Q9BTM1 | 0.43 | $1.4 \times 10^{-02}$ | $3.1 \times 10^{-02}$ | 0.86 | $4.22 \times 10^{-07}$ | $7.78 \times 10^{-06}$ | 0.60 | $2.1 \times 10^{-06}$ | $2.7 \times 10^{-05}$ |

The 2024 control group showed 20 DEPs (*P*_adj_<0.05), including 4 upregulated and 16 downregulated proteins (**Table S4**), substantially fewer than in the music-stimulated cohorts. These proteins were mainly associated with lacrimal secretion, ocular surface homeostasis, and innate immunity. Eleven of the 20 DEPs overlapped with proteins detected in the music cohorts, but several showed opposite directions of regulation. Among the 15 proteins that were consistently replicated between the 2024 and 2025 cohorts with concordant directions of effect, LACRT was the only protein also differentially expressed in the control group with the same direction of regulation. This overlap was not greater than expected by chance (Hypergeometric test, *P*(X≥1)=0.552); alternatively, this overlap may reflect the physiological role of LACRT in maintaining ocular surface homeostasis and its responsiveness to non-specific stimuli. Therefore, the 14-protein replicated signature identified in the music-stimulated cohorts was largely absent from temporal changes observed in the 2024 untreated control group, indicating that it is primarily associated with musical stimulation rather than spontaneous temporal variability.

No DEPs were detected in the 2025 control group, further supporting a music-induced proteomic response. When combining the 2024 and 2025 control groups to increase statistical power, the 14-protein replicated signature of the music-stimulated cohorts remained absent in the combined control dataset.

### Pathway analyses

Functional enrichment analyses based on GO Biological Process and Reactome identified coherent biological signatures within each cohort and a subset of pathways consistently enriched across both datasets. We first examined the top 20 most significantly enriched pathways in each cohort using the top DEPs (*P*_adj_<0.05 and |log_2_FC|≥0.5).

In the 2024 cohort (**Table S5**), GO Biological Process enrichment revealed a signature dominated by humoral immunity, hemostasis, coagulation, and lipid metabolism (**Figure 3A**). The most significant terms included humoral immune response (*P*_adj_=2.2×10^−3^), antimicrobial humoral response (*P*_adj_=2.2×10^−3^), regulation of body fluid levels (*P*_adj_=3.0×10^−3^), blood coagulation, coagulation, hemostasis (all *P*_adj_=3.0×10^−3^), and wound healing (*P*_adj_=3.5×10^−3^). Additional enriched processes involved acute-phase response, defense response to bacterium, and lipoprotein metabolism. Reactome analysis showed strong concordance, identifying pathways related to fibrin clot formation (*P*_adj_=8.3×10^−5^), platelet degranulation (*P*_adj_=9.6×10^−5^), response to elevated platelet cytosolic Ca2^+^ (*P*_adj_ =9.6×10^−5^), plasma lipoprotein assembly (*P*_adj_=1.1×10^−4^), and platelet activation, signaling and aggregation (*P*_adj_=1.4×10^−3^), supporting coordinated regulation of immune defense, coagulation, platelet biology, and lipid transport.

**Figure 3.**
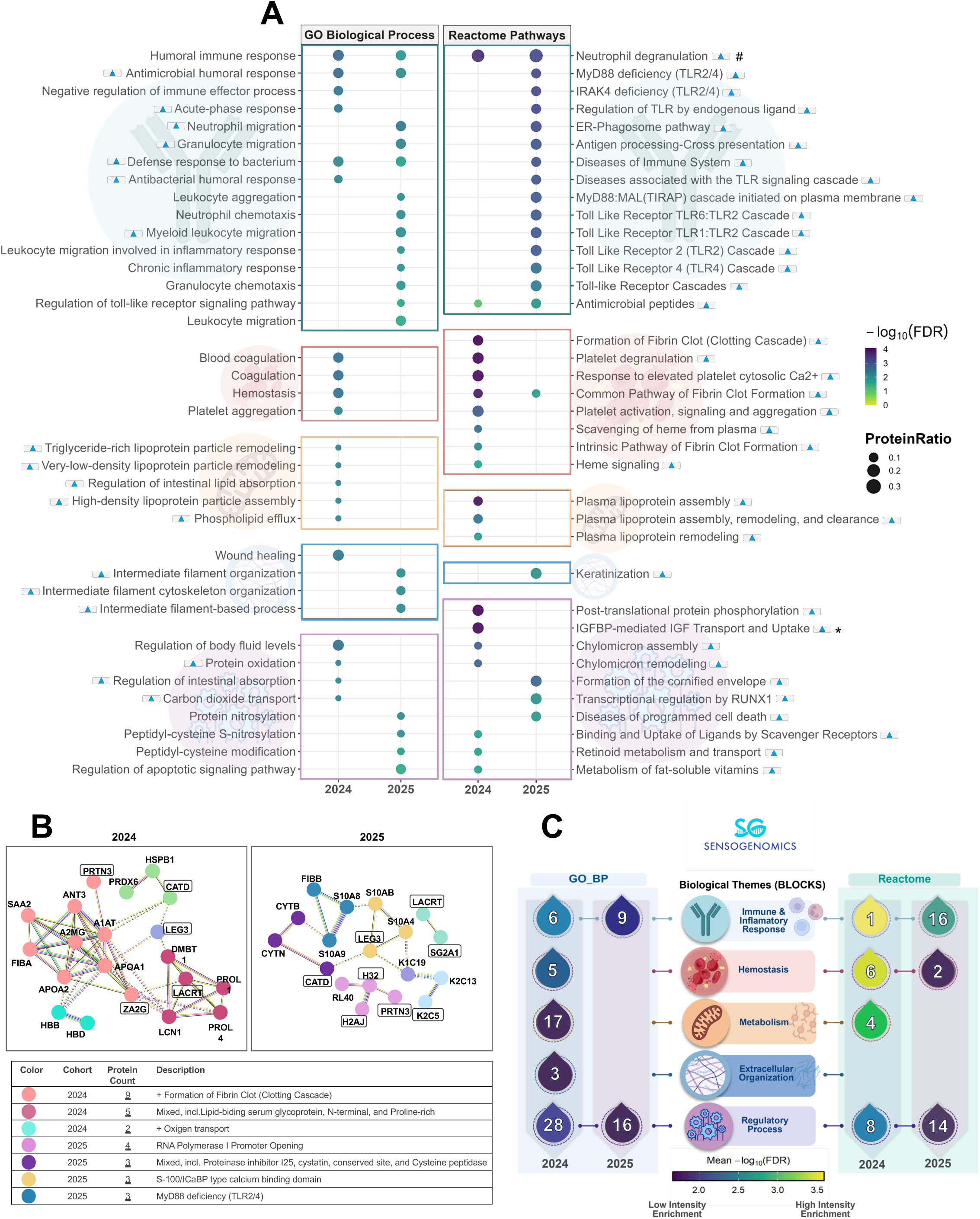
Functional enrichment analyses of the 2024 and 2025 music-stimulated cohorts based on selected differentially expressed proteins (*P*_adj_<0.05; |log_2_FC| ≥ 0.5). (A) Top 20 significantly enriched pathways identified in the 2024 and 2025 cohorts using Gene Ontology (GO) Biological Process and Reactome enrichment analyses. The complete lists of enriched pathways are provided in **Tables S4–S5**. (B) Protein–protein interaction (PPI) networks constructed from the selected DEPs identified in each cohort. Only the clusters annotated by STRING are indicated in the table below. (C) Summary of all enriched pathways identified in each cohort (**Tables S4–S5**), grouped into broader functional categories to provide an overview of the main biological processes associated with musical stimulation. For panel (A), * indicates that the complete name of the pathway is “Regulation of Insulin-like Growth Factor (IGF) transport and uptake by Insulin-like Growth Factor Binding Proteins (IGFBPs)”, which has been abbreviated here for convenience; # indicates that the pathway “neutrophil degranulation” is upregulated in the 2024 cohort but downregulated in the 2025 cohort.

The 2025 cohort (**Table S6**) displayed a signature dominated by innate immunity and inflammatory responses. GO analysis identified enrichment of neutrophil migration (*P*_adj_=3.6×10^−3^), granulocyte migration (*P*_adj_=6.6×10^−3^), antimicrobial humoral response (*P*_adj_=9.0×10^−3^), neutrophil chemotaxis (*P*_adj_=9.0×10^−3^), myeloid leukocyte migration (*P*_adj_=9.0×10^−3^), humoral immune response (*P*_adj_=1.5×10^−2^), defense response to bacterium (*P*_adj_=2.1×10^−2^), and regulation of Toll-like receptor signaling (*P*_adj_=2.7×10^−2^), together with intermediate filament organization (**Figure 3A**). Reactome analysis supported these findings, with MyD88 and IRAK4 deficiency pathways, neutrophil degranulation (all *P*_adj_=6.8×10^−4^), TLR2 and TLR4 cascades, formation of the cornified envelope, and keratinization among the most enriched pathways. Overall, the 2024 cohort was characterized by humoral immunity, coagulation, platelet activation, and lipid metabolism, whereas the 2025 cohort predominantly reflected innate immune, neutrophil-mediated, Toll-like receptor, and epithelial structural pathways.

Separate enrichment analyses showed that all significant pathways in both cohorts were driven by upregulated proteins, with no significant enrichment detected among downregulated proteins (**Figure 3A**).

PPI network analysis of the top DEPs (*P*_adj_<0.05; |log_2_FC|≥0.5) revealed modular organizations consistent with the enrichment results (**Figure 3B**). In the 2024 cohort, the main cluster was associated with coagulation and fibrin clot formation, whereas additional modules involved lipid-binding proteins, oxygen transport, and salivary proteins. In the 2025 cohort, smaller clusters corresponded to innate immune signaling, including MyD88/TLR2/4 pathways, S100 proteins, and cystatin/proteinase inhibitor families. Several replicated proteins (CATD, LEG3, PRTN3, H2AJ, H32, K2C5, SG2A1, and LACRT) occupied central network positions, supporting the biological relevance of the shared molecular signature.

To summarize the overall response, all significantly enriched GO and Reactome terms were grouped into major functional themes (**Figure 3C**). Despite cohort-specific differences, both datasets converged on five broad biological blocks: immune and inflammatory responses, hemostasis, metabolism, extracellular organization, and regulatory processes. Immune and inflammatory pathways were the dominant theme in both cohorts, whereas hemostatic and metabolic processes were more prominent in 2024 and regulatory pathways were relatively more represented in 2025.

### Protein co-expression modules analysis and functional enrichment

A consensus *WGCNA* was performed using proteomic profiles from both cohorts. After constructing signed consensus networks, three biologically relevant consensus modules were identified, comprising 54 (turquoise; hereafter SBP1 module), 25 (brown; hereafter KV320 module), and 22 (blue; hereafter MSLN module) proteins (**Figure 4A–B**; **Table S7**). Proteins assigned to the grey module were excluded from downstream analyses because they did not belong to any coherent co-expression cluster. All three modules showed significant associations with musical stimulation in both cohorts, with consistent effect directions across datasets (**Figure 4C**). The MSLN module showed the strongest negative association with music (2024: *r*=−0.57, *P*=2.48×10^−16^; 2025: *r*=−0.33, *P*=2.06×10^−4^). The SBP1 module displayed a weaker but reproducible negative correlation (2024: *r*=−0.19, *P*=0.012; 2025: *r*=−0.19, *P*=0.043), whereas the KV320 module was positively correlated with music (2024: *r*=0.23, *P*=0.003; 2025: *r*=0.47, *P*=4.99×10^−8^) (**Figure 4C**). All three modules were highly preserved between cohorts (**Figure 4D**), with *Zsummary* values of 7.94 (SBP1), 6.50 (KV320), and 5.24 (MSLN), all exceeding the threshold for moderate preservation (*Zsummary*>2).

**Figure 4.**
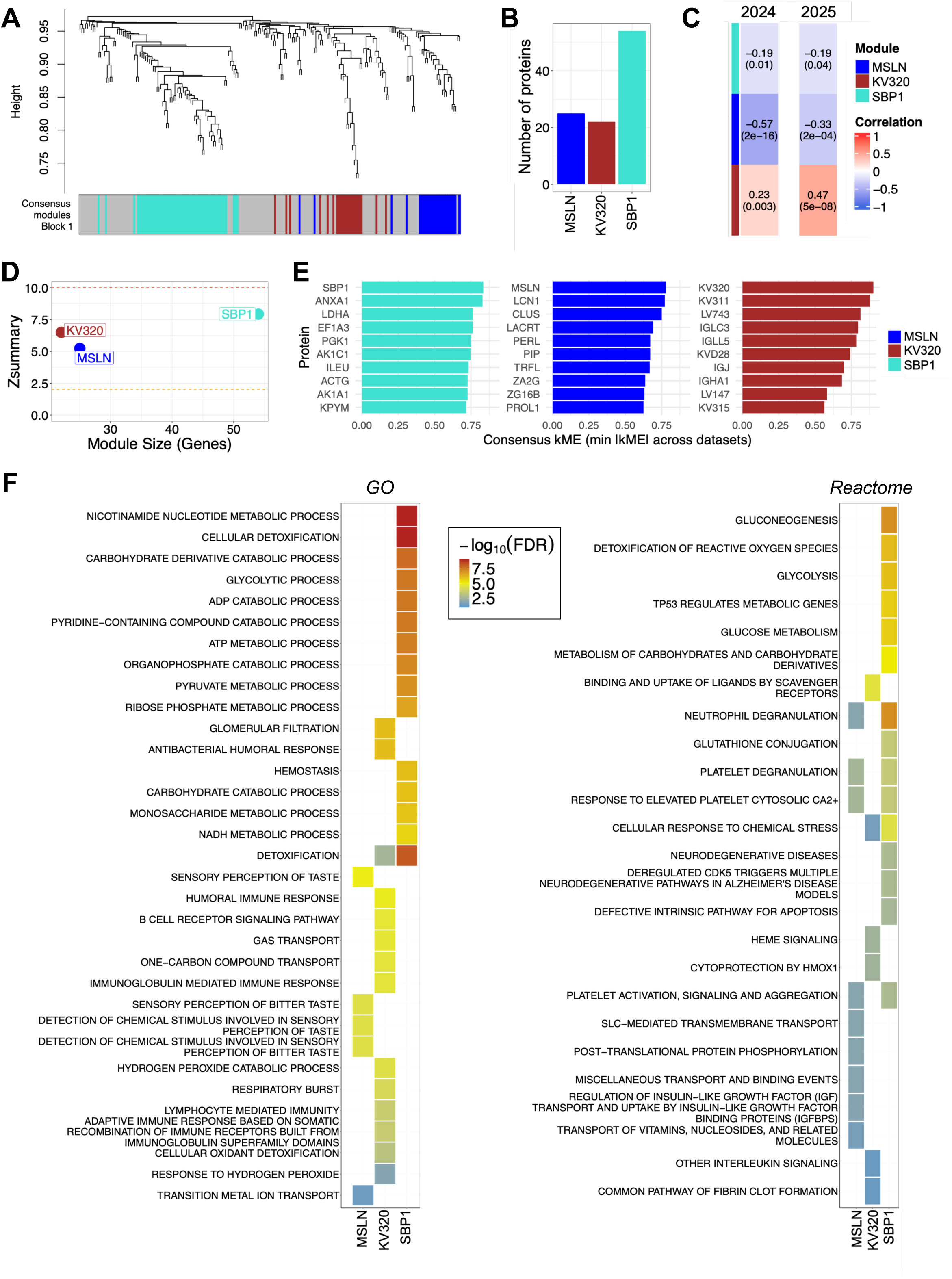
Co-expression network analysis of tear proteins in response to musical stimulation. (A) Consensus modules of co-expressed proteins were identified across the 2024 and 2025 datasets. Individual modules are represented by distinct colors. (B) Number of proteins assigned to each consensus module. (C) Heatmap showing module–trait correlations obtained from *WGCNA* in the 2024 and 2025 datasets. Correlation coefficients between each module and the musical stimulation trait (TP1 *vs*. TP2) are shown in the upper part of each cell, with the corresponding *P*-value indicated in parentheses. (D) Module preservation analysis displaying *Zsummary* values (y-axis) as a function of module size (number of proteins; x-axis). Thresholds for moderate (*Zsummary*=2) and strong (*Zsummary*=10) preservation are indicated by orange and red dashed lines, respectively. (E) Top 10 hub proteins identified within each consensus module, ranked according to their consensus module membership scores. (F) Functional enrichment analysis of modules significantly associated with musical stimulation using GO terms and Reactome pathways. Only the 15 most significantly enriched terms or pathways per module are shown.

Hub proteins were ranked using a consensus module membership score derived from both cohorts (**Figure 4E**). The SBP1 module was characterized by proteins involved in metabolism and cellular stress responses, including SBP1, ANXA1, LDHA, EF1A3, and PGK1. The MSLN module was enriched in secretory and epithelial proteins such as MSLN, LCN1, CLUS, LACRT, and LPO. The KV320 module showed the strongest hub connectivity and was almost entirely composed of immunoglobulins, with all top 10 hub proteins belonging to the immunoglobulin superfamily.

Functional characterization revealed distinct but partially overlapping biological signatures across the three modules (**Figure 4F**; **Table S8**). The SBP1 module showed the broadest repertoire, including oxidative stress responses, cellular detoxification, energy metabolism, innate immunity, coagulation, and wound healing. The MSLN module was mainly associated with sensory perception, secretory functions, and transport processes, while also sharing platelet and neutrophil activation pathways with SBP1. In contrast, the KV320 module was dominated by adaptive immune processes, particularly humoral and immunoglobulin-mediated responses, together with erythrocyte-associated functions and cytoprotective pathways.

Additional details on the co-expression modules are provided in **Supplementary Text 2**.

### Machine-learning classification based on reproducible protein signature

Of the fifteen proteins consistently replicated across the 2024 and 2025 cohorts (**Table 1**), which remained statistically significant after multiple-testing correction and exhibited concordant directions of regulation in both datasets, LACRT was excluded because it was also downregulated in the control group and was therefore not considered specific to musical stimulation. The remaining fourteen replicated, music-specific DEPs were used to develop classification models, using the 2024 cohort as the training set and the 2025 cohort as the independent test set. All four machine-learning algorithms achieved excellent discrimination between TP1 and TP2 samples in the 2024 cohort (discovery [“training”] set), with area under the ROC curve (AUC) values of 0.874 for LR, 0.942 for RF, 0.952 for SVM, and 0.873 for LDA (**Figure 5A**). The same biomarker panel also showed good classification performance in the independent 2025 cohort (replication set), yielding AUC values of 0.870 for LR, 0.812 for RF, 0.754 for SVM, and 0.860 for LDA (**Figure 5B**).

**Figure 5.**
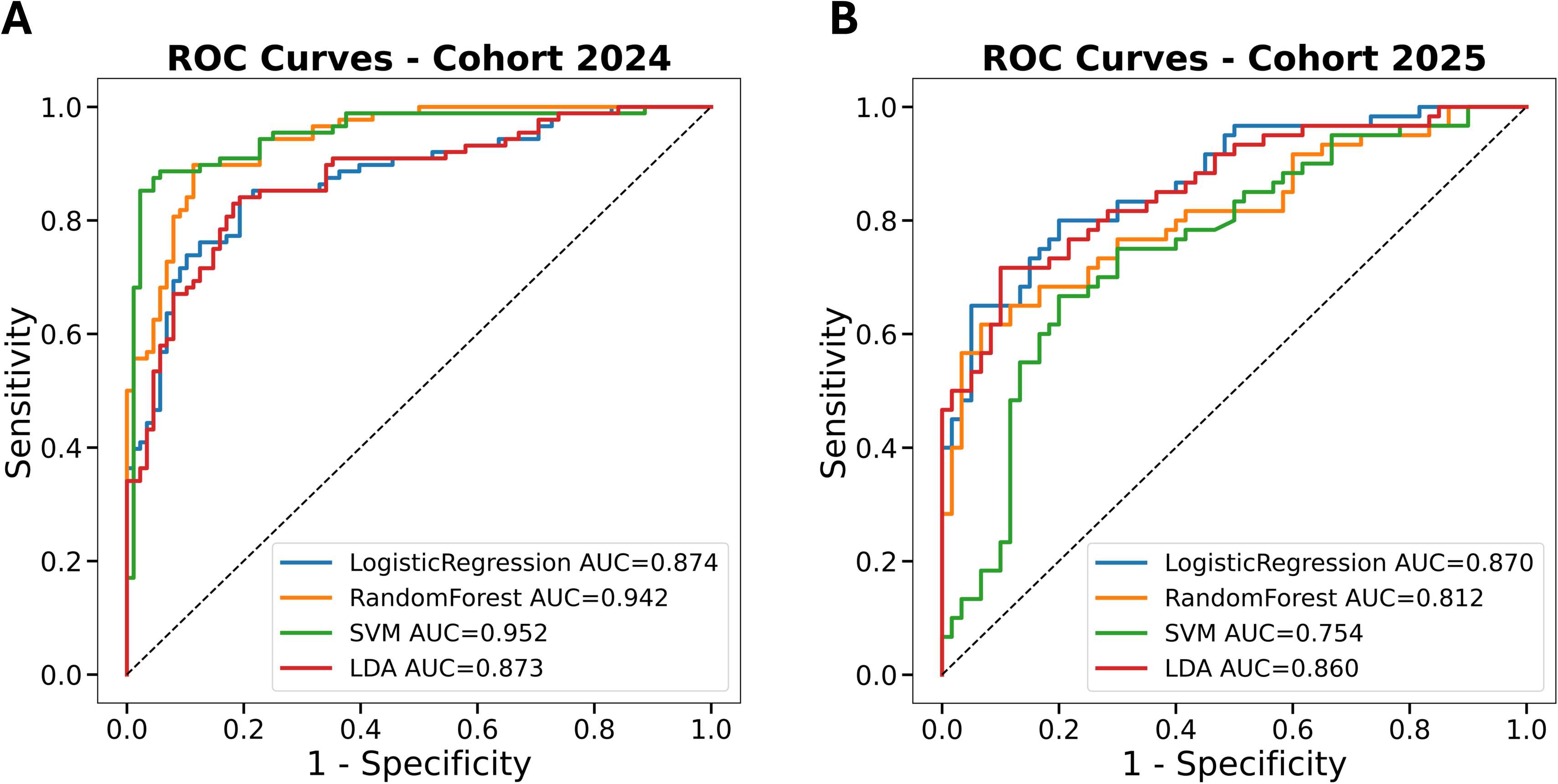
ROC-based classification performance of the 14-protein signature in the 2024 and 2025 music-stimulated cohorts. Receiver operating characteristic (ROC) curves showing the ability of the 14 reproducibly differentially expressed proteins (DEPs) to discriminate post-stimulation (TP2) from pre- stimulation (TP1) samples using four classification models: logistic regression, random forest, support vector machine, and linear discriminant analysis. Models were trained on the 2024 cohort and externally validated on the independent 2025 cohort. ROC curves are shown for the 2024 training cohort (A) and the independent 2025 validation cohort (B). The area under the ROC curve (AUC) is indicated for each model.

Finally, three of the fourteen proteins should be interpreted with caution, as their differential expression may reflect technical or biological noise rather than a true response to musical stimulation. These proteins were SG2A1, K2C5, and G3PT (see **Supplementary Text 3** for detailed discussion). We recomputed the classification models using an 11-DEP signature and found that their predictive performance remained virtually unchanged compared with the 14-protein model (**Figure S2**).

## Discussion

Music elicits complex emotional, cognitive, and physiological responses, yet the molecular mechanisms underlying these effects remain poorly understood. It is also largely unknown whether music exposure induces measurable changes in the human proteome, the primary functional machinery of biological activity. Tear fluid is an attractive biofluid for addressing this question because it reflects both local ocular processes and broader systemic physiology. The tear proteome has been extensively studied in ocular diseases, including glaucoma, dry eye disease, and diabetic retinopathy ^43, 44^, and has recently emerged as a promising surrogate matrix for investigating systemic and brain-related biomarkers, supported by the close anatomical and physiological connection between the eye and the central nervous system (CNS) ^45, 46^. Proteomic studies of tears and tear-derived extracellular vesicles have identified molecular signatures associated with neurodegenerative disorders ^47, 48^, and substantial overlap has been reported between proteins and pathways detected in tears, cerebrospinal fluid, and blood, suggesting that tears can capture biological processes occurring within the CNS ^47^. Using quantitative proteomics in two independent music-stimulated cohorts together with two music-unstimulated control groups, the present study identified reproducible protein alterations, biological pathways, and co-expression networks associated with music exposure in tears from healthy donors (**Figure 6**).

**Figure 6.**
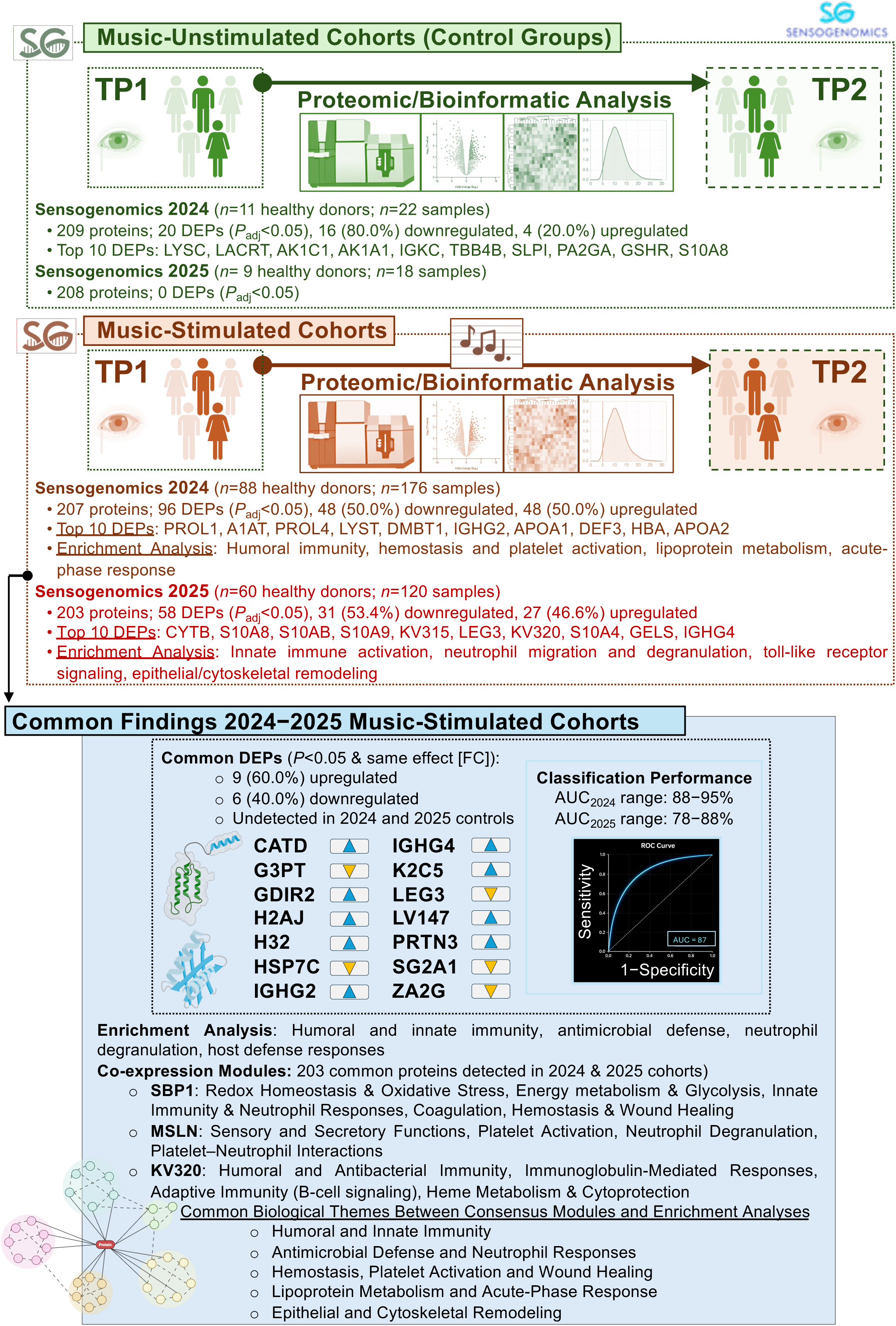
Schematic overview of the main findings of the present study across the four cohorts (2024 and 2025 music-stimulated and unstimulated control groups), highlighting the biological features shared by the two music-stimulated cohorts, including differentially expressed proteins (DEPs), enriched biological pathways, and protein co-expression modules.

The proteins consistently modulated after musical stimulation delineate a coordinated molecular response that is best interpreted within the framework of the lacrimal functional unit (LFU), a neurosecretory and neuroimmune system in which sensory input, autonomic regulation, immune homeostasis, and lacrimal gland secretion are tightly integrated ^49, 50^. Rather than representing isolated ocular changes, the observed alterations suggest that tear fluid captures downstream molecular adaptations elicited by neural activity associated with music (**Supplementary Text 3**). This interpretation is supported by the remarkable reproducibility of the proteomic signature across two independent cohorts, indicating that the tear proteome constitutes a sensitive peripheral readout of complex sensory stimulation.

A major finding was the coordinated regulation of proteins involved in neuroimmune signaling. The upregulation of immunoglobulin-related proteins (IGHG2, IGHG4, and IGLV1-47), together with modulation of LEG3, PRTN3, and GDIR2, suggests that musical stimulation induces adaptive neuroimmune responses rather than classical inflammatory activation. Immunoglobulins are increasingly recognized as active participants in neuroimmune communication beyond their canonical immune functions ^51, 52^, whereas reduced LEG3 expression is consistent with attenuation of pro-inflammatory signaling mediated by activated microglia ^53^. Increased PRTN3 and GDIR2 levels further support the coordinated regulation of innate immune pathways and Rho GTPase signaling, both of which are involved in bidirectional communication between the nervous and immune systems^54–56^. Beyond immune regulation, musical stimulation also affected proteins related to proteostasis, metabolic homeostasis, and epigenetic regulation. Increased CATD abundance is compatible with enhanced lysosomal protein turnover ^57^, whereas downregulation of HSP7C may reflect reduced activation of stress-responsive chaperone systems during physiological relaxation ^58, 59^. Modulation of ZA2G suggests systemic metabolic adaptation ^60–62^, while increased H3.2 and H2AJ raises the possibility that sensory stimulation is accompanied by transient epigenetic responses associated with neuronal plasticity ^63–65^. These coordinated molecular changes indicate that music engages an integrated physiological program linking neural activity with immune regulation, protein quality control, metabolic adaptation, and cellular homeostasis. Additional discussion is provided in **Supplementary Text 3**.

SG2A1, K2C5, and G3PT should be interpreted cautiously because their biological relevance in tear physiology remains uncertain and they may reflect technical rather than biological variation (**Supplementary Text 3**). Consequently, these proteins were considered exploratory findings requiring independent validation.

The high predictive performance of the 14-protein signature across two independent cohorts demonstrates that the coordinated molecular changes identified here are robust and sufficiently specific to discriminate biological responses to musical stimulation with high accuracy. Importantly, a reduced 11- protein version retained comparable predictive performance.

The pathway enrichment differences between cohorts likely reflect the complexity and interindividual variability of biological responses to music rather than inconsistency between datasets. Music perception involves coordinated activation of neural, endocrine, immune, and metabolic networks whose responses may vary according to individual characteristics, emotional engagement, previous musical experience, and contextual factors. Despite cohort-specific differences, both datasets converged on a common biological framework involving antimicrobial defense, innate and humoral immunity, neutrophil-associated responses, and coagulation-related pathways. GO Biological Process indicates that humoral immune response, antimicrobial humoral response, and defense response to bacterium pathways were significantly enriched in both cohorts (**Figure 3A**, **Tables S4–S5**), while Reactome analysis identified neutrophil degranulation among the most significantly enriched pathways in each dataset, together with pathways related to innate immune responses and antimicrobial defense. This convergence supports the robustness of these signatures and suggests that musical stimulation elicits coordinated systemic responses involving evolutionarily conserved pathways linked to host defense, immune regulation, and tissue homeostasis. The simultaneous involvement of immune, inflammatory, and hemostatic processes is consistent with growing evidence that music influences neuroimmune and physiological regulatory networks beyond the CNS ^66, 67^.

Consensus network analysis identified three reproducible protein co-expression modules associated with music stimulation across the independent 2024 and 2025 cohorts (**Supplementary Text 3**). All modules exhibited moderate to high preservation between datasets, indicating that the observed molecular responses are unlikely to be cohort-specific. The identification of highly connected consensus hub proteins shared between cohorts further highlights central molecular mediators of the response to music. The SBP1 module contained the largest number of proteins and exhibited the broadest functional repertoire, encompassing oxidative stress responses, energy metabolism, innate immunity, neutrophil activation, coagulation, and wound healing. The MSLN module showed the strongest negative correlation with musical stimulation and was primarily associated with secretory activity, extracellular signaling, and transmembrane transport, while also sharing platelet and neutrophil activation pathways with SBP1. In contrast, the KV320 module showed coordinated upregulation after musical stimulation and was strongly enriched for humoral immunity, immunoglobulin-mediated responses, B-cell receptor signaling, and lymphocyte-mediated immunity. Most hub proteins within this module belonged to the immunoglobulin superfamily, indicating a highly interconnected antibody-centered network. The consensus co-expression analysis provided an independent systems-level validation of the biological processes identified by conventional enrichment analyses. Both approaches converged on pathways related to humoral and antimicrobial immunity, neutrophil activation, coagulation, platelet function, inflammatory responses, and metabolic adaptation, supporting the existence of a coordinated physiological response to musical stimulation that integrates metabolic, immune, inflammatory, and hemostatic processes across independent cohorts.

Reproducible molecular signatures of musical stimulation were detected in tear fluid despite the high physiological and environmental variability of this biofluid. Unlike blood, the tear film constitutes an open and highly dynamic biological interface continuously shaped by local, systemic, and environmental factors. The identification of convergent DEPs, biological pathways, and co-expression networks across two independent cohorts therefore provides strong evidence that musical stimulation can induce biologically meaningful and reproducible molecular responses. Although individual proteins and pathways differed partially between cohorts, the consistent involvement of immune, inflammatory, host-defense, and hemostatic processes supports the existence of a common physiological response program elicited by music exposure.

The inclusion of two time-matched music-unstimulated control groups substantially strengthens the interpretation of our findings. Although a small number of proteins exhibited significant temporal changes in the absence of music, these changes were observed only in the 2024 control group, and overlap with the music-associated proteomic signature was minimal. Several shared proteins showed opposite directions of regulation, further reducing biological concordance between control and music-stimulated groups. Among the 15 proteins reproducibly differentially expressed across the two independent music cohorts, only LACRT was also differentially expressed in the 2024 control group with the same direction of regulation. The remaining 14 replicated proteins were absent from temporal changes observed in the control groups, indicating that the reproducible molecular signature associated with music exposure is largely distinct from spontaneous fluctuations in tear composition or repeated sampling effects. Consistent with this interpretation, no significant DEPs were detected in the 2025 control group. Furthermore, combining the stimulated and unstimulated cohorts to increase statistical power did not alter the results or their interpretation.

Despite these promising findings, several limitations should be considered. First, although the study was designed to minimize external stimuli unrelated to music, residual confounding cannot be completely excluded. However, the paired-sample design represents a major strength because each post-stimulation sample was compared with its corresponding baseline sample from the same individual, reducing interindividual variability and enabling assessment of within-subject proteomic changes induced by music. Second, the observed proteomic alterations require replication and further characterization in independent populations, different ecological settings, and alternative exposure durations to determine their generalizability and temporal dynamics. An additional limitation relates to the analytical depth of the proteomic workflow. Although SWATH-MS provides robust, reproducible, and highly quantitative measurements, the number of proteins that can be confidently identified and quantified is inherently constrained by sample complexity, spectral library coverage, and instrument performance. Consequently, some low-abundance proteins and subtle molecular changes may not have been detected. Future studies using complementary proteomic approaches and continued methodological improvements may further expand proteome coverage and refine the molecular characterization of the response to musical stimulation. Finally, the sampling framework was designed to capture short-term molecular responses and therefore does not permit evaluation of the long-term effects of musical stimulation on protein expression or their potential relationship with the broader physiological and health benefits attributed to music exposure ^8, 68, 69^.

## Conclusions

Short-term musical stimulation induces measurable and reproducible changes in the tear proteome of healthy individuals. Across two independent cohorts, convergent alterations were consistently detected at multiple levels, including individual proteins, biological pathways, protein co-expression networks, and predictive models. The robustness of these findings is supported by the replication of 14 proteins with concordant regulation across cohorts, the strong correlation of their effect sizes, the preservation of shared functional signatures, and the robust performance of the resulting protein panel in discriminating pre- and post-stimulation samples while remaining largely unchanged in music-unstimulated controls. Together, these observations indicate that the detected molecular signature reflects a genuine biological response to music rather than random variation or temporal fluctuations. Beyond establishing tears as a sensitive, non-invasive biofluid for monitoring physiological responses to auditory stimulation, these findings provide new insights into the coordinated modulation of immune, inflammatory, epithelial, and homeostatic processes induced by music. Future studies should determine whether similar proteomic signatures are elicited across different populations, disease conditions (e.g., Alzheimer’s disease, autism, or brain injury), musical genres, and exposure durations. From an evolutionary perspective, the reproducibility of these molecular responses raises the possibility that music engages ancient and conserved biological programs linking emotional processing with immune and homeostatic regulation.

## Acknowledgements

The authors would like to express their appreciation to the study investigators of the Sensogenomics network (sensogenomics.com; Sensogenomics Working Group [see Annex]), as well as the nursery and laboratory service at the Hospital Clínico Universitario de Santiago de Compostela, for their invaluable dedication and support. We would also like to acknowledge the Real Filharmonía de Galicia (www.rfgalicia.org), the musicians of the Banda Municipal of Santiago de Compostela, the Auditorio de Galicia for their support, and donors. This research project was made possible through the access granted by the Galician Supercomputing Center (CESGA) to its supercomputing infrastructure. The supercomputer FinisTerrae III and its permanent data storage system have been funded by the Spanish Ministry of Science and Innovation, the Galician Government, and the European Regional Development Fund (ERDF). The research of the group was supported by: *i*) GAIN IN607B 2020/08 and IN607A 2023/02, and EUTERPE_adn (Programa de Cooperación Interreg-VI POCTEP; Ref. 0313_EUTERPE_ADN_1_E) (to A.S.), IIN607A2021/05 (to F.M.-T.) and IN677D 2024/06 (to A.G.-C.), and *ii*) Consorcio Centro de Investigación Biomédica en Red de Enfermedades Respiratorias (CB21/06/00103; to A.S. and F.M.-T.). AG-C is supported by the Miguel Servet contract (CP23/00080), funded by the Instituto de Salud Carlos III (ISCIII) and co-funded by the European Union. The funders were not involved in the study design, collection, analysis, interpretation of data, the writing of this article, or the decision to submit it for publication.

## Ethics approval and consent to participate

Written informed consent was obtained from all the participants in the present study. The Ethics Committee of Xunta de Galicia approved the present project (Registration code: 2020/021), and the study was conducted in accordance with the guidelines of the Helsinki Declaration.

## Consent for publication

All participants have permitted publication of the project’s findings.

## Availability of data and materials

The mass spectrometry proteomics data have been deposited to the ProteomeXchange Consortium via the PRIDE ^70^ partner repository with the dataset identifier PXD081873.

## Author’s contribution

AS conceived the study. AS, FM-T, and LN coordinated the project. AS and FM-T secured the funding. SBB performed the proteomic analyses. AS, CMC, and LN coordinated participant recruitment and sample collection. ACM, AGC, AS, MCT, and NEZM performed the data analyses and prepared the first draft of the manuscript. All authors contributed to study coordination, sample collection, and data acquisition. All authors critically revised the manuscript and approved the final version.

## Legend to the supplementary figures and tables

**Table S1. Demographic characteristics of the music-stimulated cohorts and music-unexposed control groups.** Age is presented as range and mean age, and sex as the number of females and males. Pairwise comparisons between groups were performed using the Mann–Whitney U test for age and Fisher’s exact test for sex distribution.

**Table S2. Differentially expressed proteins identified between pre- (TP1) and post-stimulation (TP2) samples in the 2024 cohort.** *Protein*, *Gene*, and *UniProt* indicate the protein name, gene symbol, and UniProt accession number, respectively. *AveExpr* represents the average expression level, log_2_FC the log_2_ fold change (TP2 *vs*. TP1), *B* the log-odds of differential expression, *t* the moderated *t*- statistic, *P* the nominal significance, and *P*_adj_ the Benjamini–Hochberg false discovery rate (FDR)-adjusted significance.

**Table S3. Differentially expressed proteins identified between pre- (TP1) and post-stimulation (TP2) samples in the 2025 cohort.** *Protein*, *Gene*, and *UniProt* indicate the protein name, gene symbol, and UniProt accession number, respectively. *AveExpr* represents the average expression level, log_2_FC the log_2_ fold change (TP2 *vs*. TP1), *B* the log-odds of differential expression, *t* the moderated *t*- statistic, *P*-value the nominal significance, and *P*_adj_ the Benjamini–Hochberg false discovery rate (FDR)-adjusted significance.

**Table S4. Differentially expressed proteins identified between pre- (TP1) and post-stimulation (TP2) samples in the control group.** *Protein*, *Gene*, and *UniProt* indicate the protein name, gene symbol, and UniProt accession number, respectively. *AveExpr* represents the average expression level, log_2_FC the log_2_ fold change (TP2 *vs*. TP1), *B* the log-odds of differential expression, *t* the moderated *t*- statistic, *P*-value the nominal significance, and *P*_adj_ the Benjamini–Hochberg false discovery rate (FDR)-adjusted significance.

**Table S5. Gene Ontology (GO) Biological Process and Reactome enrichment analyses based on differentially expressed proteins (DEPs) meeting the criteria of *P*<0.05 and |log_2_FC|<0.5 identified in the 2024 cohort after musical stimulation.** *ProteinRatio* denotes the ratio of proteins in the input dataset mapped to each enriched term, and *BgRatio* the corresponding ratio in the background set. *P*-value indicates the enrichment significance, while *P*_adj_ and *Q-value* correspond to multiple testing-adjusted significance values. *geneID* lists the proteins contributing to the enrichment, and *Count* indicates the number of proteins associated with each term.

**Table S6. Gene Ontology (GO) Biological Process and Reactome enrichment analyses based on differentially expressed proteins (DEPs) meeting the criteria of *P*<0.05 and |log_2_FC|<0.5 identified in the 2025 cohort after musical stimulation.** *ProteinRatio* denotes the ratio of proteins in the input dataset mapped to each enriched term, and *BgRatio* the corresponding ratio in the background set. *P*-value indicates the enrichment significance, while *P*_adj_ and *Q-value* correspond to multiple testing-adjusted significance values. *geneID* lists the proteins contributing to the enrichment, and *Count* indicates the number of proteins associated with each term.

**Table S7. Modules significantly associated with musical stimulation in tear samples.** The left table reports module–trait correlations between TP1 and TP2, including correlation coefficients, corresponding *P*-values, and module preservation (*Zsummary*) scores. The right table lists the top 10 hub proteins within each significant module, ranked according to consensus module membership. Module membership values for the 2024 and 2025 datasets, together with consensus module membership scores, are provided for each protein.

**Table S8. Functional enrichment analysis of tear protein co-expression modules significantly associated with musical stimulation.** Enrichment results were obtained using GO Biological Process and Reactome databases, with significantly enriched biological processes and pathways reported for each module.

**Figure S1**. Scatter plots showing the correlation of log_2_FC values between the music-stimulated cohorts and the corresponding unstimulated control groups for the 2024 and 2025 cohorts.

**Figure S2**. ROC-based classification performance of the 11-protein signature in the 2024 and 2025 cohorts and the musical-unstimulated control group. Receiver operating characteristic (ROC) curves showing the ability of the 14 reproducible DEPs to discriminate post- (TP2) from pre-stimulation (TP1) samples using four classification models: logistic regression, random forest, support vector machine, and linear discriminant analysis. Models were trained using the 2024 cohort and externally validated on the independent 2025 cohort. ROC curves are shown for the 2024 training cohort (A) and the independent 2025 validation cohort (B). The area under the ROC curve (AUC) is indicated for each model.

## Sensogenomics Working Group

Antonio Salas Ellacuriaga – PI; Federico Martinón-Torres – PI; Laura Navarro Ramón – Coordinator

*GenPoB/GenVip - Instituto de Investigación Sanitaria (IDIS) (alphabetic order)*

Alba Camino Mera, Albert Padín Villar, Alberto Gómez Carballa, Alejandro Pérez López, Alicia Carballal Fernández, Ana Cotovad Bellas, Ana Isabel Dacosta Urbieta, Ana María Pastoriza Mourelle, Ana María Senín Ferreiro, Andrés Muy Pérez, Antía Rivas Oural, Antonio Justicia Grande, Antonio Piñeiro García, Anxela Cristina Delgado García, Belén Mosquera Pérez, Blanca Díaz Esteban, Carlos Durán Suárez, Carmen Curros Novo, Carmen Gómez Vieites, Carmen Rodríguez-Tenreiro Sánchez, Celia Varela Pájaro, Claudia Navarro Gonzalo, Cristina Serén Trasorras, Cristina Talavero González, Einés Monteagudo Vilavedra, Estefanía Rey Campos, Esther Montero Campos, Fernando Álvez González, Fernando Caamaño Viñas, Francisco García Iglesias, Gloria Viz Rodríguez, Hugo Alberto Tovar Velasco, Irene Álvarez Rodríguez, Irene García Zuazola, Irene Rivero Calle, Iria Afonso Carrasco, Isabel Ferreirós Vidal, Isabel Lista García, Isabel Rego Lijo, Iván Prieto Gómez, Iván Quintana Cepedal, Jacobo Pardo Seco, Jesús Eirís Puñal, José Gómez Rial, José Manuel Fernández García, José María Martinón Martínez, Julia Cela Mosquera, Julia García Currás, Julián Montoto Louzao, Lara Martínez Martínez, Laura Navarro Marrón, Lidia Piñeiro Rodríguez, Lorenzo Redondo Collazo, Lúa Castelo Martínez, Lucía Company Arciniegas, Luis Crego Rodríguez, Luisa García Vicente, Manuel Vázquez Donsión, María Dolores Martínez García, María Elena Gamborino Caramés, María Elena Sobrino Fernández, María José Currás Tuala, María Martínez Leis, María Soledad Vilas Iglesias, María Sol Rodriguez Calvo, María Teresa Autran García, Marina Casas Pérez, Marta Aldonza Torres, Marta Bouzón Alejandro, Marta Lendoiro Fuentes, Miriam Ben García, Miriam Cebey López, Montserrat López Franco, Narmeen Mallah, Natalia García Sánchez, Natalia Vieito Perez, Nour El Zahraa Mallah, Patricia Regueiro Casuso, Ricardo Suárez Camacho, Rita García Fernández, Rita Varela Estévez, Rosaura Picáns Leis, Ruth Barral Arca, Sandra Carnota Antonio, Sandra Viz Lasheras, Sara Pischedda, Sara Rey Vázquez, Sonia Marcos Alonso, Sonia Serén Fernández, Susana Rey García, Vanesa Álvarez Iglesias, Victoria Redondo Cervantes, Vanesa Álvarez Iglesias, Wiktor Dominik Nowak, Xabier Bello Paderne, Xabier Mazaira López

*Nursing volunteers (alphabetical order)*

Alejandra Fernández Méndez, Ana Isabel Abadín Campaña, Ana María León Caamaño, Ana María Buide Illobre, Ángeles Mera Cores, Carmen Nieves Vastro, Carolina Suarez Crego, Concepción Rey Iglesias, Cristina Candal Regueira, Dolores Barreiro Puente, Elvira Rodríguez Rodríguez, Eugenia González Budiño, Eva Rey Álvarez, Fernando Rodríguez Gerpe, Gemma Albela Silva, Isabel Castro Pérez, Isabel Domínguez Ríos, José Ángel Fernández de la Iglesia, José Cruces Vázquez, José Luis Cambeiro Quintela, José Ramón Magariños Iglesias, Julia Rey Brandariz, Julio Abel Fernández López, Luisa García Vicente, Manuel González Lito, Manuel González Lijó, Manuela Pérez Rivas, Margarita Turnes Paredes, María Aurora Méndez López, María Begoña Tomé Arufe, María Campos Torres, María del Carmen Baloira Nogueira, María del Carmen García juan, María Esther Moricosa García, María Luz Chao Jarel, María Martínez Leis, María Mercedes Jiménez Santos, María Salomé Buide Illobre, María Victoria López Pereira, Mercedes Jorge González, Mercedes Isolina Rodríguez Rodríguez, Miren Payo Puente, Natalia Carter Domínguez, Olga María Reyes González, Pilar Mera Rodríguez, Purificación Sebio Brandariz, Salomé Quintáns lago, Yolanda Rodríguez Taboada, María Pereira Grau.

*Other volunteers (alphabetical order)*

Alba Arias Gómez, Alejandro Moreno Díaz, Ana Arca Marán, Astro González Guirado, Brais García Iglesias, Carlos Sánchez Rubín, Carmen Otero de Andrés, Clara Pérez Errazquin Barrera, Claudia Rey Posse, Cristina Rojas García, Eduardo Xavier Giménez Bargiela, Elena Gloria Morales García, Fabio Izquierdo García Escribano, Gabriel Guisande García, Jaime López Martín, Lara Pais Ramiro, Lucía Rico Montero, Luís Estévez Martínez, Manuel Estévez Casal, María Aránzazu Palomino Caño, María Rubio Valdés, Marisol Nogales Benítez, Miryam Tilve Pérez, Nuria Villar Muiños, Pablo Del Cerro Rodríguez, Pablo Pozuelo Martínez Cardeñoso, Salma Ouahabi El Ouahabi, Santiago Vázquez Calvache

