## Supplementary figures and images for "A prospective multi-cohort study identifies reproducible molecular responses to musical stimulation in the human tear proteome"

### Figure S1

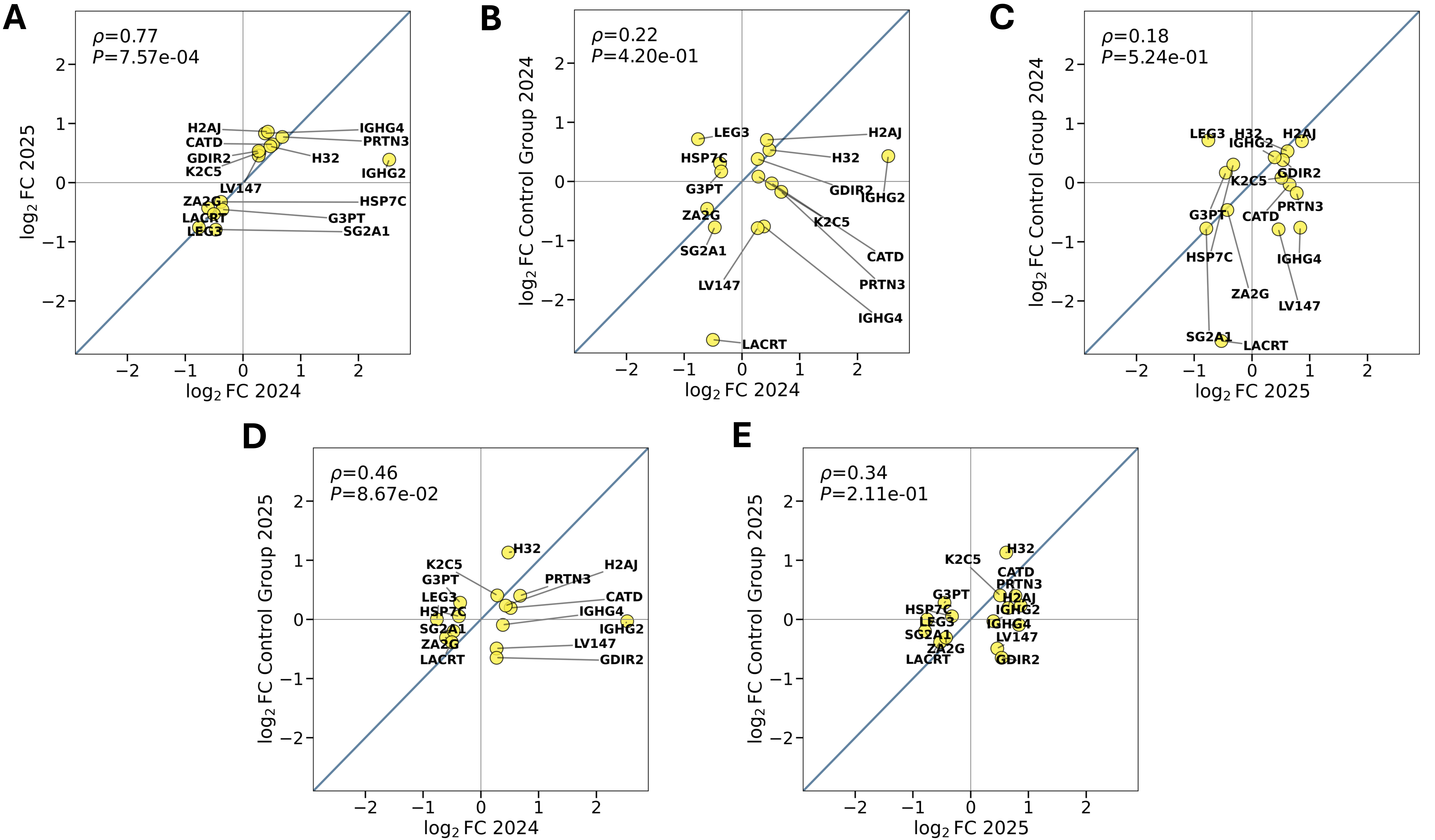

### Figure S2

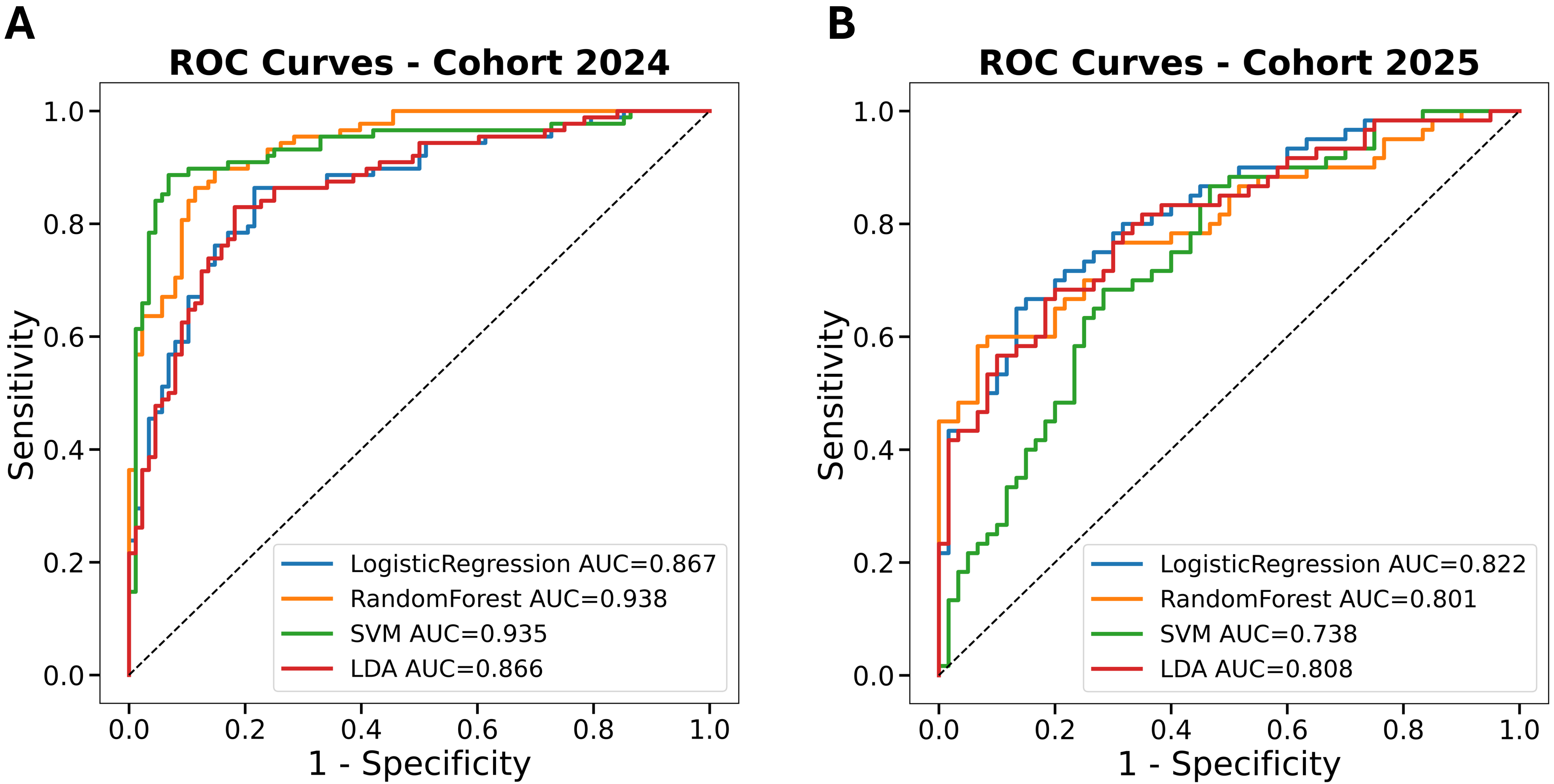
