## Supplementary Text 1 for "A prospective multi-cohort study identifies reproducible molecular responses to musical stimulation in the human tear proteome"

**Supplementary Text 1 – Extended Methods**

Experimental design and sampling

Within the framework of the Sensogenomics project (http://sensogenomics.com) ^1^, a prospective, controlled, repeated-measures experimental study was conducted, comprising two independent music-stimulated cohorts and two concurrent non-stimulated control cohorts.

The two music-stimulated cohorts were prospectively designed as an independent discovery cohort (2024) and an independent replication cohort (2025) to assess the reproducibility of the molecular response to musical stimulation. Both experimental cohorts were based on independent 50-minute live classical music concerts held at the Auditorio de Galicia (Santiago de Compostela, Spain). The first concert (discovery cohort; cohort 2024) took place on September 27, 2024, and the second (replication cohort; cohort 2025) on October 3, 2025, under comparable experimental conditions. Both events were organized in collaboration with the Real Filharmonía de Galicia (https://www.rfgalicia.org/es/) and the Banda Municipal de Música de Santiago de Compostela. To minimize potential sources of bias, participants were not informed of the musical repertoire before the concerts. Throughout the performances, they remained seated and engaged exclusively in passive listening, without physical activity or social interaction. The concert environment was intentionally maintained as comfortable, calm, and non-invasive to reduce stress and other external factors that might influence physiological responses. Both cohorts followed an identical experimental design, in which the musical program was structured to sequentially evoke two basic emotional states through works with contrasting acoustic and expressive characteristics. In both cohorts, the first half of the concert was designed to elicit sadness, whereas the second half was intended to evoke joy.

The repertoire was selected to maximize contrasts in tempo, rhythmic activity, melodic profile, and orchestration while preserving the ecological validity of a live symphonic performance. Sensogenoma24 was conducted by Baldur Brönnimann and performed by the Real Filharmonía de Galicia together with the Banda Municipal de Música de Santiago de Compostela. The sadness section included works by Michael Nyman, Ennio Morricone, John Williams, Samuel Barber, and John Barry, characterized predominantly by slower tempi, sustained melodic lines, lower rhythmic density, and reduced metric activity. The joy section comprised works by Aram Khachaturian, Leonard Bernstein, John Philip Sousa, Paul Dukas, Johann Strauss II, and John Williams, featuring faster tempi, greater rhythmic regularity, higher metric activity, and brighter orchestration. Likewise, Sensogenoma25 followed the same experimental framework but featured a predominantly classical repertoire. Conducted by Sebastian Zinca and Casiano Mouriño, the sadness section included works by Tchaikovsky, Brahms, James Horner, Puccini, and Elgar, whereas the joy section comprised compositions by Kabalevsky, Dvořák, Prokofiev, Khachaturian, and Bizet. Although the specific repertoire differed between cohorts, both concerts were designed to maximize the emotional and acoustic contrast between the sadness and joy conditions while maintaining comparable experimental settings.

Concurrently with the musically stimulated groups, two independent control groups comprising 11 and 9 healthy donors, respectively, were monitored at the Auditorio de Galicia under the same temporal schedule and environmental conditions as the music-stimulated participants. Instead of attending the concert, these individuals remained in a separate sound-isolated room within the same venue, where they were not stimulated to the musical performance or any other structured sensory stimulation. Participants spent the session in a relaxed and informal environment, engaging in spontaneous conversation while remaining otherwise inactive. These contemporaneous unstimulated control cohorts were included to account for molecular changes associated with the passage of time, sample collection procedures, and the experimental setting itself. Their inclusion enabled the discrimination of proteomic alterations specifically associated with musical stimulation from spontaneous temporal fluctuations unrelated to the intervention, thereby strengthening the specificity of the molecular signature identified in the music-stimulated cohorts.

Under clinical supervision, tear samples were collected from all participants using Schirmer strips without anesthesia at two time points (TPs): immediately before (TP1) and after (TP2) the concerts or the corresponding control session. The strips were placed under the lower eyelid for a maximum of 5 min and were removed earlier if 15 mm of wetting was achieved within 2-3 min. Samples that reached 15 mm in less than 1 min were discarded to avoid contamination from reflex tearing. Conversely, strips that did not reach 15 mm after the full 5-min collection period were considered indicative of dry eye and were excluded from further analyses. No evidence of conjunctival degeneration or skin contamination was observed in the analyzed samples, as proteins typically associated with skin tissue were not detected. The collected strips were placed in 1.5 mL Eppendorf tubes and stored at -80 °C for further processing. Participants diagnosed with dry eye disease and/or other ocular pathologies were excluded from tear donation. Furthermore, all participants were instructed in advance to avoid wearing makeup on the day of sampling.

All participants enrolled in this study were healthy donors. Cohort 2024 consisted of 88 participants (54 females and 34 males), aged 20-87 years (mean age = 51 years). Cohort 2025 included 60 participants (36 females and 24 males), aged 18-83 years (mean age = 50 years). The control cohorts comprised 11 and 9 participants, respectively (2024 control: 9 females and 2 males, aged 25-69 years, mean age = 52 years; 2025 control: 8 females and 1 male, aged 44-65 years, mean age = 54 years). No significant differences in sex distribution were observed between the 2024 and 2025 cohorts (Fisher’s exact test, *P*=0.866; OR=1.06), indicating comparable demographic characteristics across the two experimental cohorts. No significant differences in sex distribution were observed between the control cohorts and either the 2024 cohort (Fisher’s exact test, *P*=0.319; OR=0.35) or the 2025 cohort (*P*=0.307; OR=0.33). Similarly, age distributions did not differ significantly between the 2024 and 2025 cohorts (Mann-Whitney U test, *P*=0.690). Comparison of age distributions showed no statistically significant difference between the 2024 control cohort and the 2024 cohort (Mann-Whitney U test, *P*=0.057), whereas the 2024 control cohort was modestly older than the 2025 cohort (*P*=0.034). Despite this modest age difference, the groups showed comparable demographic characteristics overall, with similar mean ages (51, 50, and 52 years for the 2024, 2025, and 2024 control cohorts, respectively) and no significant differences in sex distribution.

For proteomic analyses, pooled samples from the musically stimulated participants and the control participants were initially generated and used to construct an in-house spectral library, maximizing protein identification and proteome coverage. Quantitative analyses were subsequently performed on individual samples to compare protein abundance between TP1 and TP2.

**Figure 1** illustrates the experimental design of the study.

Protein quantification by SWATH

*Creation of the mass spectral (MS) library*

To build the MS/MS spectral libraries, the peptide solutions were analyzed using a shotgun data-dependent acquisition (DDA) approach with micro-LC-MS/MS. To obtain a representative set of peptides and proteins across all samples, pooled vials of samples from each group were prepared using an equal mixture of the original samples. One 4 mL of each pool was separated into a micro-LC system Ekspert nLC425 (Eksigen, Dublin, CA, USA) using a YMC-Triart C18 column (150μm×0.3mm, 12 mm, s-3µm) (YMC CO.Japan) at a flow rate of 5µL/min. Water and ACN, both containing 0.1% formic acid, were used as solvents A and B, respectively. The gradient run consisted of 5% to 95% B for 30 min, 5 min at 90% B, and finally 5 min at 5% B for column equilibration, for a total run time of 40 min. As the peptides eluted, they were directly injected into a hybrid quadrupole-TOF mass spectrometer TripleTOF® 6600 system (AB SCIEX) operated with a data-dependent acquisition system in positive ion mode. A Micro source (SCIEX) was used as the interface between microLC and MS, with an applied voltage of 2600 V. The acquisition mode consisted of a 250 ms survey MS scan from 400 to 1250 m/z followed by an MS/MS scan from 100 to 1500 m/z (25 ms acquisition time) of the top 65 precursor ions from the survey scan, for a total cycle time of 2.8 s. The fragmented precursors were then added to a dynamic exclusion list for 15 s; any singly charged ions were excluded from the MS/MS analysis.

Peptide and protein identification was performed using ProteinPilot software (v5.0.1, SCIEX). MS/MS data were searched against the Human Uniprot database (<https://www.uniprot.org/>), with iodoacetamide specified as the cysteine alkylating agent. A false discovery rate (FDR) threshold of 1 was applied at both peptide and protein levels. The MS/MS spectra of the confidently identified peptides were subsequently used to generate a spectral library for Sequential Window Acquisition of All Theoretical Mass-Spectra (SWATH) analysis. SWATH peak extraction was performed using the MS/MSALL with SWATH Acquisition MicroApp (v2.0, SCIEX) integrated into PeakView software (v2.2, SCIEX). Only peptides identified with a confidence score above 99%, as determined by the ProteinPilot database search, were incorporated into the spectral library.

*Relative quantification by SWATH acquisition*

The objective of this study was to investigate the effect of auditory stimuli on the identification and quantification of tear proteins in healthy individuals, with high confidence (>99%), using TripleTOF MS in a SWATH-MS quantification mode. The SWATH-MS data-independent acquisition (DIA) approach enabled comprehensive and reproducible profiling of tear proteins across samples collected before and after music exposure. By systematically acquiring fragment ion data across predefined mass windows, SWATH-MS provided broad proteome coverage and consistent quantification of protein abundance changes associated with the auditory intervention. Furthermore, the permanent digital record generated by SWATH-MS allowed retrospective interrogation of the dataset, thereby enhancing confidence in protein identification and enabling detailed assessment of proteomic responses to music exposure. The high sensitivity, mass accuracy, and acquisition speed of the TripleTOF platform further supported reliable protein detection and quantification, while minimizing missing values and improving reproducibility across biological samples.

Briefly, individual tear samples (4 *μ*L) were analyzed using the same LC-MS instrumentation and chromatographic gradient employed for spectral library generation. The SWATH acquisition method consisted of repeated cycles comprising an initial TOF MS survey scan (400–1500 m/z, 50 ms acquisition time), followed by 100 TOF MS/MS scans acquired in high-sensitivity mode (400–1500 m/z, 50 ms acquisition time) across sequential overlapping precursor isolation windows of variable width (1 m/z overlap) covering the 400–1250 m/z mass range. The total cycle time was 6.3 s. For each sample set, the width of the 100 variable SWATH windows was optimized according to precursor ion density derived from the DDA analyses using the SCIEX SWATH Variable Window Calculator.

*Quantitative data analysis*

Targeted extraction of fragment-ion chromatographic traces from the SWATH datasets was performed in *PeakView* (v2.2) using the *SWATH Acquisition* *MicroApp* (v2.0). Data processing was carried out using the spectral library generated by the DDA (shotgun) analyses. Up to ten peptides per protein and seven fragment ions per peptide were selected based on signal intensity; shared and modified peptides were excluded. Ion chromatograms were extracted using a 5-min retention time window and a mass tolerance of 30 ppm. SWATH quantitation was performed for all proteins included in the spectral library that had been identified by *ProteinPilot* at an FDR<1%. The retention times from the peptides that were selected for each protein were realigned in each run according to the retention time (iRT) peptides for some proteins present in all samples and eluted along the whole time axis. Extracted ion chromatograms were generated for each selected fragment ion, and peptide abundances were calculated by summing the corresponding fragment-ion peak areas. *PeakView* assigned a score and estimated FDR for each peptide according to chromatographic and spectral features; only peptides with an FDR<1% were retained for protein quantification. Protein abundances were calculated by summing the peak areas of the corresponding peptides.

Integrated peak areas (processed.mrkvw files generated in *PeakView*) were directly exported to the *MarkerView* software (SCIEX) for relative quantitative analysis. This export generated three datasets containing quantitative information at the fragment-ion, peptide, and protein levels. *MarkerView* has been widely used in SWATH-MS studies because of its robust data-independent quantitation framework ^2-5^. The software employs algorithms that accurately detect chromatographic and spectral features directly from the raw SWATH data. Data alignment compensates for minor variations in both mass accuracy and retention time, ensuring reliable comparison of identical analytes across samples. To account for potential variability introduced during sample preparation, global normalization was performed using the total summed peak area of all detected peptides and transitions within each sample.

Weighted protein co-expression network analysis

Protein co-expression networks were constructed using the Weighted Gene Co-expression Network Analysis approach implemented in the *WGCNA* R package ^6^. To identify robust protein networks reproducibly associated with the response to music stimulation, a consensus network analysis was performed using the 2024 and 2025 cohorts. The analysis was restricted to the 203 proteins common in both cohorts and available at the two sampling TPs. Consensus networks were generated from normalized protein abundance data and corrected for repeated measures using the *removeBatchEffect* function from the *limma* package ^7^. The soft-thresholding power was evaluated independently in each cohort. A soft-thresholding power of β=5 was selected for downstream signed weighted network construction, as this value produced the best-balanced approximation to a scale-free network topology in both datasets.

Consensus modules were identified with a minimum module size of 20 proteins, a module splitting sensitivity parameter of *deepSplit*=2, and a module merging threshold of 0.20. Network topology was summarized using topological overlap matrices (TOMs), and module structure was visualized through hierarchical clustering dendrograms. For each consensus module, module eigengenes (MEs) were calculated separately in both cohorts. Associations and statistical significance between module eigengenes and music stimulation status (TP1 and TP2) were assessed independently in each cohort.

The reproducibility of consensus modules was evaluated using a module preservation analysis implemented in the *WGCNA* package. Module preservation statistics were estimated using 1,000 permutation tests. Preservation was assessed using the composite *Zsummary statistic metric*. As a reference, *Zsummary* values >10, between 2 and 10, and <2 indicate strong preservation, moderate preservation, and lack of preservation, respectively.

Hub proteins, defined as the most representative proteins within each module, were identified within each consensus module using module membership, calculated independently in both cohorts. Modules were named according to their hub proteins. To identify hub proteins that were reproducibly connected to the same module in both datasets, a consensus module membership score was calculated as the minimum absolute kME observed across cohorts:

kME_consensus_=*min*(|kME_2024_|,|kME_2025_|)

Proteins showing opposite signs of module membership between cohorts were excluded from hub ranking. Within each module, proteins were ranked according to their consensus kME values.

Functional pathway analysis of the significant modules was performed through the *clusterProfiler* ^8^ R package, using FDR for multiple test correction, and *P*-value and *Q*-value thresholds set both to 0.05. We interrogated GO and Reactome as the reference database, with a maximum gene-set size of 550 and a minimum gene-set size of 20.
