## Supplementary Text 2 for "A prospective multi-cohort study identifies reproducible molecular responses to musical stimulation in the human tear proteome"

**Supplementary Text 2 – Extended Results**

**Detailed characterization of consensus co-expression modules**

The hub proteins, the most representative proteins within each module, were ranked according to a consensus module membership score derived from both cohorts (**Figure 4E**). Within the SBP1 module, the highest-ranked hub proteins included SBP1, ANXA1, LDHA, EF1A3, and PGK1, all showing consensus kME values >0.75 (**Figure 4E**). The top hub protein (SBP1; KMEconsensus = 0.83) is the methanethiol oxidase encoded by the SELENBP1 gene. The MSLN module was characterized by hub proteins such as MSLN, LCN1, CLUS, LACRT, and LPO, with consensus kME values ranging from 0.67 to 0.78 (**Figure 4E**). The top hub protein of the MSLN module (MSLN; KMEconsensus = 0.77) is mesothelin encoded by the MSLN gene. The KV320 module displayed the strongest hub connectivity overall, with KV320, KV311, LV743, IGLC3, and IGLL5 showing consensus kME values > 0.78 and reaching a maximum value of 0.90 (**Figure 4E**). These five proteins belong to the immunoglobulin superfamily, specifically to the lambda and kappa subunits. Interestingly, all top 10 hub proteins in this module are also immunoglobulins.

Functional assessment of the significantly correlated modules revealed several associated biological pathways (**Figure 4F**; **Table S7**). The SBP1 module showed the highest number of significantly enriched processes (*n* = 81 for GO Biological Process and *n* = 42 for Reactome; **Table S7**). Results from GO Biological Process revealed a strong enrichment for terms related to cellular detoxification, redox homeostasis, energy metabolism, and innate immune functions. The most significantly enriched terms were cellular detoxification and nicotinamide nucleotide metabolic process (*P_adj_* = 1.03 × 10^−9^; **Table S7**, **Figure 4F**). Consistently, GO Biological Process categories indicating activation of antioxidant mechanisms and metabolic adaptation to oxidative stress were significantly enriched, including cell redox homeostasis, response to oxidative stress, detoxification of reactive oxygen species, and cellular response to oxidative and chemical stress (**Table S7**, **Figure 4F**). A second major functional cluster was related to glucose metabolism, carbohydrate metabolism, and bioenergetic processes. Highly enriched pathways included glycolytic process, pyruvate metabolic process, ATP metabolic process, NADH metabolic process, nicotinamide nucleotide metabolic process, carbohydrate catabolic process, monosaccharide metabolic process, and ribose phosphate metabolic process (**Table S7**, **Figure 4F**). In addition to metabolic pathways, the SBP1 module showed significant enrichment for hemostatic and wound-healing functions, including hemostasis, blood coagulation, coagulation, and wound healing (**Table S7**, **Figure 4F**). Finally, immune and inflammatory pathways were also among the functional clusters defined by this module, including chemotaxis, granulocyte chemotaxis, granulocyte migration, and interleukin-1 production (**Table S7**, **Figure 4F**).

Reactome pathway analysis strongly corroborated the enrichment results from GO Biological Process. The most significantly enriched pathway was neutrophil degranulation (*P_adj_* = 9.49 × 10^−8^), highlighting proteins involved in innate immune activation and granule-mediated antimicrobial responses (**Table S7**, **Figure 4F**). The overlap between GO Biological Process and Reactome results also points to a coordinated metabolic program centered on glucose utilization and energy production, including gluconeogenesis, glycolysis, glucose metabolism, and metabolism of carbohydrates and carbohydrate derivatives (**Table S7**, **Figure 4F**). Reactome further identified significant enrichment of detoxification of reactive oxygen species and oxidative stress responses, consistent with the involvement of this module in antioxidant processes detected in GO Biological Process analysis. Finally, Reactome analysis also highlighted pathways associated with platelet activation and degranulation, consistent with GO Biological Process enrichment detected for coagulation and wound-healing processes (**Table S7**, **Figure 4F**).

Functional analysis of the MSLN module with GO Biological Process revealed an enrichment profile dominated by sensory perception and secretory processes. The most significant biological process was sensory perception of taste (*P_adj_* = 1.20 × 10^−5^; **Table S7**, **Figure 4F**). Closely related terms, including detection of chemical stimulus involved in sensory perception of bitter taste and sensory perception of bitter taste, were also significantly associated. Additional enrichment of transition metal ion transport suggested involvement in metal-binding and transport functions (**Table S7**, **Figure 4F**). Reactome analysis supported this secretory function, highlighting SLC-mediated transmembrane transport, transport of vitamins, nucleosides, and related molecules, and miscellaneous transport and binding events. Furthermore, enrichment of pathways related to platelet degranulation, response to elevated platelet cytosolic Ca^2+^, and neutrophil degranulation was detected, overlapping with significant processes involving the SBP1 module, which suggests an inter-module functional cross-talk in processes related to platelet-neutrophil interactions (**Table S7**, **Figure 4F**).

Finally, GO Biological Process analysis of the proteins within the KV320 module identified antibacterial humoral response and glomerular filtration as the most significantly enriched terms (*P_adj_* = 8.71 × 10^−7^; **Table S7**, **Figure 4F**). Additional enrichment of humoral immune response, immunoglobulin-mediated immune response, B-cell receptor signaling pathway, and lymphocyte-mediated immunity further demonstrated an important adaptive immune involvement of this module (**Table S7**, **Figure 4F**). Remarkably, nestled directly alongside these immune pathways was a significant signature for erythrocyte-associated physiological processes, including gas transport and one-carbon compound transport. Functional enrichment of the KV320 module using Reactome revealed pathways associated with scavenger receptor-mediated ligand uptake, heme signaling, HMOX1-dependent cytoprotection, and cellular responses to chemical stress. It is also important to note that Reactome enrichment results were based only on seven genes of the KV320 module due to the lack of Entrez annotation for some of the immunoglobulin proteins (**Table S7**, **Figure 4F**).
