## Supplementary Text 3 for "A prospective multi-cohort study identifies reproducible molecular responses to musical stimulation in the human tear proteome"

**Supplementary Text 3 – Extended Discussion**

**Replicated DEPs in 2024 and 2025 cohorts**

The present study provides evidence that musical stimulation induces reproducible changes in the human tear proteome, revealing that tear fluid can capture molecular adaptations elicited by complex sensory experiences. Rather than representing an isolated ocular phenomenon, these changes are most plausibly interpreted within the framework of the lacrimal functional unit (LFU), a highly integrated neurosecretory and neuroimmune system in which sensory input, autonomic output, immune regulation, and lacrimal gland secretion are tightly coordinated. Because the LFU is continuously regulated by central and peripheral neural circuits, alterations in tear composition may represent downstream molecular signatures of neural activity elicited by music. This physiological framework provides a mechanistic basis for interpreting the coordinated molecular changes observed. Under this framework, the proteins identified converge into interconnected biological processes involving neuroimmune signalling, proteostasis, metabolic regulation, and cellular adaptation.

Neuroimmune signalling within the LFU

One of the most prominent biological themes emerging from the proteomic analysis was the modulation of proteins involved in neuroimmune signalling. Among these, three immunoglobulin-related proteins (IGHG2, IGHG4, and LV147) were consistently upregulated following musical stimulation, reinforcing accumulating evidence that immunoglobulins participate in neural physiology beyond their classical role in adaptive immunity. Thus, Immunoglobulin Heavy Constant Gamma 2 (IGHG2) is of special interest because accumulating evidence indicates that immunoglobulin genes are expressed not only by immune cells but also by neurons and microglia, where they contribute to specialized neuroimmune communication independently of peripheral immune responses ^1^. Furthermore, the widespread distribution of neuron-derived IgG and its receptors throughout both the central and peripheral nervous systems supports an active role for immunoglobulins in bidirectional neuroimmune crosstalk ^2^. The association between IGHG2 and neurodegenerative disorders further reinforces this interpretation. Increased levels of IgG2 constant-region fragments have previously been detected in the frontal cortex of patients with advanced Alzheimer’s disease, where they are thought to reflect disturbances in immune homeostasis accompanying disease progression ^3^. Although the analyzed cohorts represent healthy individuals, the increased abundance of IGHG2 in tear fluid following musical stimulation is more likely to reflect activation of physiological neuroimmune pathways rather than pathological inflammation. The LFU is tightly regulated by trigeminal sensory afferents together with sympathetic and parasympathetic innervation, which collectively coordinate tear secretion and immune homeostasis ^4, 5, 6, 7, 8^. Consequently, music-induced neuronal activity may indirectly modulate the secretion of immune-associated proteins into the tear film. The concurrent upregulation of IGHG4 and LV147 provides additional support for this hypothesis. Although secretory IgA constitutes the predominant immunoglobulin class in tears, IgG subclasses also contribute to mucosal immune surveillance ^8^. Among them, IGHG4 is noteworthy because of its well-established immunoregulatory and anti-inflammatory properties ^9^, whereas the increased abundance of both IGHG2 and LV147 may reflect subtle remodeling of the local humoral immune repertoire following sensory stimulation. These concerted alterations indicate that musical stimulation influences neuroimmune signalling at the ocular surface, promoting adaptive changes in the tear immunoproteome that extend beyond conventional inflammatory responses.

Additional evidence supporting neuroimmune modulation comes from the coordinated regulation of LEG3 (Galectin-3), PRTN3 (Proteinase 3), and GDIR2 (Rho GDP-dissociation inhibitor 2), three proteins that participate in complementary aspects of innate immune signalling and inflammatory regulation. Among them, the downregulation of LEG3 is interesting because Galectin-3 acts as a microglia-derived lectin that is rapidly induced following central nervous system (CNS) injury, where it functions as a damage-associated molecular pattern (alarmin) capable of amplifying neuroinflammatory signalling through receptors such as TLR4 and TREM2, thereby promoting neurotoxic microglial activation associated with neurodegenerative disorders ^10, 11, 12^. Experimental studies have further demonstrated that pharmacological inhibition of Galectin-3 attenuates pro-inflammatory cytokine release while promoting neuronal survival ^11^. Consequently, the reduction in LEG3 observed after musical stimulation is consistent with a shift toward a more balanced, homeostatic neuroimmune state characterized by reduced inflammatory tone. Within the context of the lacrimal functional unit, this decrease may therefore represent a peripheral indicator of attenuated neuroimmune activation mediated through autonomic and sensory pathways linking the ocular surface to the CNS ^4, 5, 6^. Interestingly, PRTN3 exhibited the opposite pattern of regulation, being significantly upregulated following musical stimulation. Proteinase 3 is a neutrophil serine protease stored in azurophilic granules together with neutrophil elastase and cathepsin G, where it contributes to pathogen elimination and tissue remodeling during innate immune responses ^13,^ ^14^. Beyond these classical functions, recent transcriptomic analyses have implicated PRTN3 in CNS biology. In Alzheimer’s disease, PRTN3 was identified within a synaptic gene-enriched module and displayed a strong inverse correlation with amyloid pathology, suggesting that higher expression may be associated with protection against amyloid accumulation, whereas reduced levels accompany synaptic degeneration ^15^. Although the physiological significance of increased PRTN3 in tears remains to be established, its coordinated regulation together with the immunoglobulin proteins and LEG3 supports the hypothesis that musical stimulation modulates neuroimmune pathways extending beyond the ocular surface itself.

Further evidence for coordinated neuroimmune adaptation is provided by the upregulation of GDIR2. As a key regulator of Rho GTPase signalling, GDIR2 controls cytoskeletal dynamics, leukocyte migration, immune-cell activation, and inflammatory signalling ^16, 17^. Because Rho GTPases occupy a central position in the bidirectional communication between the nervous and immune systems, whereby neuronal activity influences immune responses through specialized neuroimmune circuits ^18^, increased GDIR2 expression is compatible with adaptive regulation of immune-cell function induced by sensory stimulation. The coordinated modulation of immunoglobulin proteins, LEG3, PRTN3, and GDIR2 provides convergent evidence that musical stimulation engages physiological neuroimmune pathways, highlighting the tear proteome as a sensitive peripheral readout of neural activity and immune regulation.

Proteostasis, metabolic adaptation and epigenetic regulation

Beyond neuroimmune signalling, the proteins modulated by musical stimulation converged on biological pathways involved in protein quality control, cellular stress adaptation, metabolic homeostasis, and epigenetic regulation. Thus, the molecular response to music is not restricted to immune modulation but extends to fundamental mechanisms that preserve cellular integrity and maintain physiological homeostasis.

One of the most biologically relevant changes within this functional group was the upregulation of CATD (Cathepsin D), the principal lysosomal aspartyl protease responsible for degrading proteins implicated in neurodegenerative disorders, including tau, α-synuclein, and β-amyloid ^19, 20^. By facilitating lysosomal degradation and protein turnover, CATD plays a central role in maintaining proteostasis, whereas its deficiency promotes protein aggregation, glial activation, and progressive neurodegeneration ^21^. Importantly, because the increase observed occurred in healthy individuals in the absence of pathological stimuli, it is unlikely to reflect tissue damage. Instead, the enhanced abundance of CATD is more consistent with a physiological activation of lysosomal clearance pathways and cellular quality-control mechanisms that contribute to maintaining protein homeostasis under conditions of increased sensory stimulation ^22^.

A complementary perspective on proteostasis is provided by HSP7C (Heat Shock Cognate 71-kDa Protein), a constitutively expressed member of the HSP70 family that occupies a central position within the cellular chaperone network. HSP7C facilitates protein folding, prevents protein aggregation, and participates in chaperone-mediated autophagy while simultaneously contributing to the regulation of both innate and adaptive immune responses ^23, 24^. In contrast to CATD, HSP7C was downregulated following musical stimulation. Although the biological significance of this decrease remains uncertain, it may reflect reduced activation of stress-responsive chaperone systems under conditions of physiological relaxation rather than impaired proteostasis. Because molecular chaperones are dynamically regulated according to cellular protein-folding demand, the simultaneous upregulation of lysosomal degradation pathways together with reduced expression of stress-associated chaperones may indicate a coordinated reorganization of protein quality-control mechanisms induced by musical stimulation rather than a generalized stress response.

In addition to proteins involved in proteostasis, musical stimulation also modulated proteins associated with systemic metabolic regulation. Among these, ZA2G (Zinc Alpha-2 Glycoprotein; AZGP1) was significantly downregulated. AZGP1 is a multifunctional adipokine involved in lipid mobilization, glucose homeostasis, insulin sensitivity, and immune regulation ^25, 26, 27, 28^. Under pathological conditions such as obesity and metabolic syndrome, AZGP1 expression is typically suppressed by chronic low-grade inflammation through the action of pro-inflammatory cytokines ^29, 30^. However, the reduction observed most likely represents a physiological adaptation rather than pathological dysregulation. Given the established role of AZGP1 in promoting lipolysis and energy-substrate mobilization, one plausible explanation is that music-induced relaxation decreases the metabolic demand for lipid mobilization. Although the molecular mechanisms responsible for this response remain to be elucidated, modulation of ZA2G may constitute part of the broader systemic metabolic adaptations accompanying relaxation-associated physiological states ^28, 30^.

Evidence for additional layers of regulation is provided by the increased abundance of Histone H3.2 and the histone variant H2AJ following musical stimulation. Although histones are traditionally regarded as structural components of chromatin, they are now recognized as active regulators of transcription, inflammatory signalling, neuronal plasticity, and cellular responses to environmental stimuli ^31, 32^. H2AJ has been implicated in the maintenance of persistent inflammatory signalling during cellular senescence ^33^. While the biological significance of extracellular histones in tear fluid remains largely unexplored, the coordinated increase of H3.2 and H2AJ raises the intriguing possibility that epigenetic regulatory mechanisms contribute to the molecular adaptations elicited by sensory stimulation. Although these observations remain speculative, they are consistent with an emerging body of evidence indicating that environmental stimuli, including music, can induce transient epigenetic responses associated with neuronal plasticity and long-term physiological adaptation.

The coordinated regulation of CATD, HSP7C, ZA2G, and histone proteins indicates that musical stimulation influences biological processes extending well beyond immune signalling alone. The simultaneous modulation of lysosomal protein degradation, molecular chaperone activity, metabolic homeostasis, and epigenetic regulation supports the concept that sensory stimulation promotes an integrated adaptive response aimed at preserving cellular homeostasis while accommodating changes in neural activity. Rather than representing isolated molecular events, these proteins collectively point toward a coordinated physiological program linking neural stimulation with systemic mechanisms of cellular maintenance and adaptation.

Proteins requiring cautious interpretation, and lacritin

Although most of the DEPs identified in this study could be interpreted within coherent biological pathways, three proteins, SG2A1 (Mammaglobin-B), K2C5 (Keratin type II cytoskeletal 5), and G3PT (testis-specific glyceraldehyde-3-phosphate dehydrogenase), should be interpreted with caution, as their biological relevance in the context of musical stimulation remains uncertain.

SG2A1 belongs to the secretoglobin superfamily, whose members participate in epithelial homeostasis, innate immunity, and the regulation of inflammatory responses ^34, 35, 36^. However, its specific biological function at the ocular surface has not yet been established, precluding a mechanistic interpretation of its downregulation following musical stimulation. Likewise, K2C5 warrants careful consideration because Schirmer strips recover not only tear fluid but also superficial epithelial cells. As a result, keratins are commonly detected in mass spectrometry-based tear proteomics ^37, 38^, and changes in their abundance may reflect differences in epithelial cell recovery during sample collection rather than true biological changes in the tear proteome. Furthermore, its high sequence homology with the ubiquitously expressed GAPDH increases the possibility of peptide misassignment during LC–MS/MS analysis, thereby limiting confidence in the biological interpretation of its reduced abundance. Although these proteins reached statistical significance, they should be regarded as exploratory findings pending independent validation.

LACRT (lacritin) deserves separate consideration because, despite being replicated across both experimental cohorts, it was also identified as a DEP in the untreated control group and was therefore excluded from the final panel of biomarkers specific to musical stimulation. This distinction is important given the central physiological role of LACRT in maintaining ocular surface homeostasis. Secreted primarily by the lacrimal gland, LACRT promotes basal tear secretion, epithelial renewal, and tear-film stability, and its deficiency has consistently been associated with dry eye disease, supporting its utility as a biomarker of lacrimal gland function and ocular surface health ^39, 40^. Because all participants were healthy and showed no clinical evidence of ocular surface disease, the observed reduction in LACRT abundance is unlikely to represent a pathological alteration. Instead, it is more plausibly interpreted as a transient physiological response associated with sensory stimulation. Importantly, quantitative proteomics measures relative rather than absolute protein abundance. Consequently, the apparent decrease in LACRT may reflect changes in tear volume resulting from autonomic nervous system activity, producing a dilution effect, or a transient reorganization of the lacrimal secretory program induced by sensory and emotional processing ^5, 37^. The fact that a similar reduction was observed in the control group indicates that this response is not specific to musical stimulation and reinforces the importance of including appropriate controls when identifying stimulus-specific molecular signatures.

Concluding remarks

The proteins identified in this study delineate a coordinated molecular response that extends well beyond the CNS and encompasses interconnected pathways involved in neuroimmune communication, inflammatory regulation, proteostasis, cellular stress adaptation, metabolic homeostasis, and lacrimal gland physiology. Rather than representing isolated molecular events, the reproducible modulation of these proteins across two independent cohorts supports the existence of an integrated physiological program linking sensory stimulation with neural, immune, autonomic, and metabolic regulation.

Although the precise mechanisms underlying these responses remain to be elucidated, the remarkable reproducibility of the proteomic signature across independent cohorts, together with its ability to accurately discriminate samples obtained before and after musical stimulation, provides compelling evidence that tear fluid captures systemic biological adaptations elicited by sensory experience. These findings not only identify a robust panel of candidate biomarkers of exposure to musical stimulation but also establish the tear proteome as a promising and non-invasive platform for investigating the molecular consequences of complex sensory and emotional stimuli. Future studies integrating tear proteomics with neurophysiological, transcriptomic, and metabolomic approaches will be essential to define the cellular origin of these proteins, elucidate the signalling pathways involved, and determine whether similar molecular signatures characterize other forms of sensory stimulation or neurological disorders.

**Biological interpretation of consensus co-expression modules**

Among the negatively correlated modules detected, the SBP1 module contained the largest number of proteins. The negative correlation with musical stimulation indicates a coordinated reduction in the abundance of proteins belonging to this module following stimulation. Despite showing a discrete correlation value, this module displayed the most robust functional enrichment profile. The convergence of GO and Reactome analyses suggests that these proteins participate in a coordinated adaptive response to cellular stress integrating energy metabolism, oxidative defense, and inflammatory regulation. The simultaneous enrichment of glucose metabolism and antioxidant pathways is consistent with the tight coupling between metabolic processes and redox regulation ^41^.

SBP1 emerged as a central network node of the module. This protein, encoded by the SELENBP1 gene, catalyzes the oxidation of methanethiol and is involved in redox regulation, sulfur-containing compound metabolism, intra-Golgi protein transport, and cellular differentiation ^42^. SELENBP1 has been linked to oxidative stress, inflammation, and neuronal injury, processes increasingly recognized as important regulators of brain plasticity and neural responses to environmental stimuli (REF). Overexpression of SELENBP1 has been reported in several neurological and neuropsychiatric disorders, including schizophrenia, epilepsy, multiple sclerosis, and spinal cord injury, suggesting a broader role in nervous system function and pathology ^43, 44, 45, 46, 47, 48^.

Moreover, enrichment of innate immune, neutrophil-associated, and hemostatic pathways was detected for the SBP1 module, overlapping with platelet activation and neutrophil-related processes identified in the MSLN module, which exhibited the strongest negative correlation with musical stimulation. This overlap suggests functional interactions between both modules, reflecting coordinated attenuation of cellular stress, redox imbalance, and inflammatory responses. These findings are consistent with experimental evidence showing that music exposure can prevent stress-induced and anxiety-like behaviors through restoration of hypothalamic-pituitary-adrenal axis function and reduction of oxidative stress and inflammation^49, 50, 51^.

The MSLN module exhibited the strongest negative correlation with musical stimulation and was functionally related to secretory activity, extracellular signaling, and transmembrane transport, suggesting a coordinated decrease in proteins involved in intercellular communication and regulation of circulating proteins. Interestingly, several hub proteins within this module (LCN1, LACRT, and AZGP1) were also validated as DEPs in the independent validation cohort, providing additional evidence of the central role of this module in the local response to musical stimulation.

Finally, the KV320 module showed coordinated up-regulation after musical stimulation and was enriched for pathways related to humoral immune response, immunoglobulin-mediated immunity, B-cell receptor signaling, and lymphocyte-mediated immunity, demonstrating a strong adaptive immune component, particularly involving antibody-mediated responses. Although the individual Reactome terms appeared heterogeneous, they were largely driven by the recurrent involvement of HBA1, HBA2, PRTN3, and JCHAIN, suggesting the existence of a common underlying biological process. Previous studies have reported associations between music exposure and modulation of immune parameters, including changes in immunoglobulin levels, lymphocyte activity, and cytokine production, supporting the hypothesis that musical stimulation may influence immune regulation beyond its effects on psychological well-being ^52, 53^.

The consensus co-expression network analysis also provided an independent systems-level validation of the biological processes identified by conventional enrichment analyses. The SBP1 module integrated pathways related to redox homeostasis, energy metabolism, innate immunity, neutrophil function, and hemostatic responses, thereby bridging the principal biological signatures identified independently in the 2024 and 2025 cohorts. Similarly, the MSLN module reinforced the existence of coordinated platelet-neutrophil interactions, whereas the KV320 module identified a highly interconnected immunoglobulin-centered network consistent with the enrichment of humoral and antibacterial immune responses. Beyond confirming the enrichment analyses, the network approach revealed that these biological processes are not activated independently but rather constitute tightly interconnected functional programs involving metabolic adaptation, oxidative stress regulation, innate immunity, adaptive humoral responses, and immuno-hemostatic interactions.
